# Diverse ancestral myosin motors generate and segregate distinct types of nanocluster-rich domains at the plasma membrane

**DOI:** 10.1101/2025.09.30.679578

**Authors:** Parijat Sil, Thomas S. van Zanten, Sowmya Jahnavi, Ajay Bansal, Suvrajit Saha, Bhagyashri Mahajan, Mukesh Kumar, Parvinder Pal Singh, Madan Rao, Satyajit Mayor

## Abstract

Molecular organization of the plasma membrane (PM) at nano and micron scales is critical for its function in all living cells. This emerges not only from the self-assembly of lipids and proteins but also from active forces originating in the underlying cytoskeletal cortex. These forces drive membrane molecules into non-equilibrium steady state patterns such as nanoclusters. However, the molecular agents connecting membrane organization with cytoskeletal dynamics and stresses have remained unknown. Here we show that two classes of ubiquitous ancestral non-muscle myosins are deployed for the organization of different types of membrane components. Inner-leaflet localized Class I myosins link outer-leaflet GPI-anchored molecules to juxta-membrane actin-filaments, whereas the more cortically-localized Class II myosins operate on transmembrane proteins endowed with actin-binding capacity. Consistent with an active Flory Huggins theory for phase separation, these observations show that the distinct motor-driven membrane molecules generate spatially segregated mesoscale domains, enriched in nanoclusters derived from different myosin classes. Moreover, chemically reversible post-translational modifications such as palmitoylation enable concatenation of these domains by enhancing affinity of the membrane domain constituents for each other. We anticipate that the segregation potential of the ATP-fueled cell membrane is made available for the crucial purpose of modulating information transduction because it can be regulated in space and time during the construction of signaling cascades, underpinning functional plasma membrane organization.

## Introduction

The PM of every living cell is a two-dimensional (2D) fluid bilayer of lipids and proteins whose composition is regulated in a cell type specific manner(*1*). It exhibits functional lateral heterogeneity and transverse lipid and protein asymmetry(*2–5*). Its organization is crucial for its functioning as a dynamic interface for communication with the extracellular milieu and responding to internal states of the cell (*6*, *7*). While protein asymmetry is due to the presence of distinct peripheral proteins at the inner and outer leaflet of the PM, transverse lipid asymmetry is established by ATP-driven flippases(*1*). The generation of lateral heterogeneity has been proposed to occur by interactions of membrane constituents as outlined in the fluid mosaic and the lipid raft models(*8*, *9*). However, evidence for segregation of membrane lipids at physiological temperatures, consistent with thermodynamical determined phases has been difficult to obtain(*10*–*12*). Instead, crosslinking of membrane protein(*12*, *13*) and actin-dependent clustering mechanisms (*14*, *15*) result in building membrane domains with distinct chemical compositions and properties(*12*, *16*).

Based on observations in living cells, it has been proposed(*14*) and validated *in vitro*(*17*) that actin filaments and motor activity create contractile flows culminating in dynamic platforms that promote transient nanoclustering. These contractile platforms exhibit mesoscale organization, termed *active emulsions*(*16*). Evidence for these have primarily come from observations of the dynamics and distribution of nanoclusters of outer-leaflet localized GPI-anchored proteins (GPI-APs), glycolipids and inner leaflet lipid-anchored molecules (*15*, *18*). It is expected that the characteristics of these mesoscale domains depend on the specific molecules that are assembled in the contractile platforms. In the case of GPI-APs, nanoclustering requires enrichment of cholesterol and immobilization of inner leaflet phosphatidylserine (PS), as well as trans-bilayer interactions of long acyl chain-containing lipid species at the outer and inner leaflet(*19*). Nanoclusters exhibit local lipid order (*lo),* well above the phase transition temperature (Tm) of the membranes (*19*), and recruit other *lo*-preferring components. As a consequence the mesoscale domains of GPI-APs also exhibit *lo*-like characteristics(*16*). Although *active emulsions* of GPI-APs resemble phase-separated *lo* domains, the mechanism that generates these domains is distinct from equilibrium phase separation and is likely to have different properties. These features have propelled a new understanding of the nano and mesoscale organization of the membrane for all eukaryotic cells in terms of an active composite, taking into account interactions arising from the specific composition of the membrane(*16*, *17*).

Notwithstanding the progress made in understanding the active membrane composite(*3*, *20*), the precise molecular agents that drive this activity in the cell are still unknown. We provide evidence that the ancestral and ubiquitous class I and II non-muscle myosin protein families(*21*) are the engines that power this organization. Class I and II myosins perform a diverse range of tasks in the cell. Class I facilitates endocytosis, phagocytosis, vesicle and protein transport on filamentous actin including on protrusions such as microvilli and cilia, and initiate left-right asymmetry at the organismal scale due to their chiral character, the non-muscle class II myosins are required for building contractile structures such as stress fibers and the actin cortex(*22–25*). Here we discover altogether new functions for these two classes of myosin families, in the active and regulated organization of PM components, first by studying their distribution in the cell, and then the consequence of their selective perturbation, consistent with a theoretical framework that emphasizes the role of active stresses in organizing membrane components.

In all metazoa, the actin cortex is infused with class II myosins, and here we show that these motors are necessary for the construction of nanoclusters of membrane proteins that are associated with cortical actin. Simultaneously, we find that the ubiquitously present lipid-associated class I myosins drive the nano and mesoscale arrangement of lipidic nanoclusters at the outer leaflet likely due to their ability to associate with acidic lipid species at the inner leaflet of the PM(*19*). Thus, class I and II myosins serve as the main motors at the PM and in the actin cortex, driving the nanoscale clusters of membrane lipids and proteins, respectively. Finally, we show that these motor-driven nanoclusters assemble into distinct domains with higher order organization. They exhibit two kinds of active emulsions at the mesoscale with the capacity to sort or concatenate membrane components of different chemistry, consistent with an active Flory Huggins (AFH) theory, recently proposed by us(*26*). These domains are used in regulating the function of the membrane in many cellular processes, including in cell spreading.

### Class I and Class II myosins exhibit stratified distribution at the cortex

Non-muscle myosin class I motors have a lipid-binding domain and class II motors are localized in the actin cortex, juxtaposed to the PM(*27*). Therefore, we examined the localization of these two classes of myosin motors with respect to the PM and the actin cortex. For this we imaged a representative of both class I (GFP-Myo1C) and class II myosin (mCherry-Myo2A) together with PM and actin in CHO cells (Fig 1). GFP-Myo1C colocalizes with the membrane marker in both the basal and equatorial plane, well separated from the major peak for actin (Fig. 1A-C). mCherry-Myo2A, on the other hand, is predominantly localized within the membrane-proximal actin and is enriched on thick actin stress fibers (Fig. 1A-C). Quantification of the distances involved, shows the layering of the different proteins with respect to the membrane and actin-cortex; class I myosin is distinctly membrane associated (<30 nm away) whereas class II myosin is enriched between 100-120 nm in the juxta-membrane actin cortex (Fig. 1C-E).

**Fig. 1.**
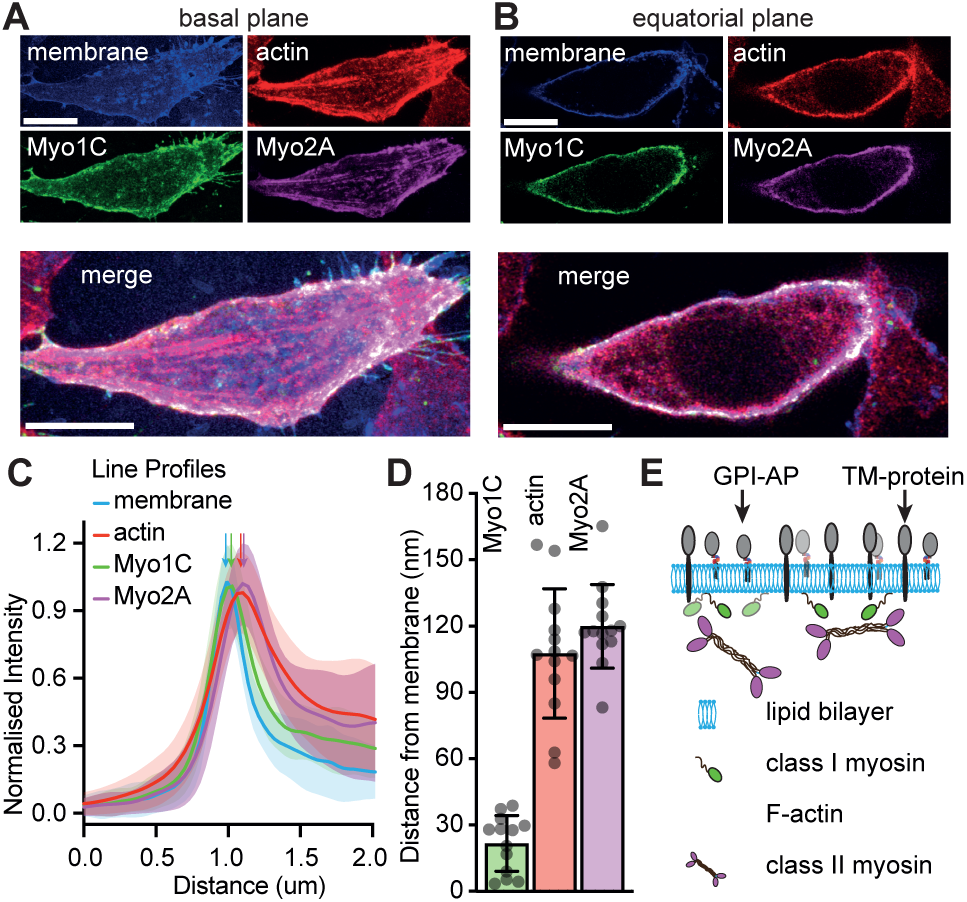
Distinct myosins decorate different layers of the cortical-actin supported plasma membrane. A,B) Representative airy scan microscopy image of a fluorescently labeled membrane marker, myosin 1C, F-actin and myosin 2A in a single cell spread on fibronectin-coated glass, focused at the cell-glass interface; a, or away from the glass; **B,C)** Plot shows the average of 69 membrane tangential line-profiles (solid lines, shaded regions depict s.d) taken from 13 cells, where the value of each line-profile starting 1 μm outside and ending 1 μm into the cell was normalized to its maximum value. **D)** Bar graphs show the average distance (±s.d) of the myosins and actin from the plasma membrane determined from the position of the fitted maximum intensity of each protein with that of the plasma membrane marker. Each point corresponds to mean distance of Myo1C, F-actin and Myo2A from the plasma membrane obtained from the measurements within a single cell; when averaged across all cells the distance corresponds to 21.7±13 nm, 107.6±29 nm, and 119.9±19 nm, respectively. **E)** Schematic shows the organization of GPI-APs at the outer leaflet of the plasma membrane, TM proteins inserted in the plasma membrane with class I myosin proteins at the inner membrane leaflet and class II myosin entangled in the cortical F-actin meshwork. Scale bars,10 μm.

### Class II myosins drive nanoclustering of transmembrane actin-binding proteins

Since the cortex forms a shell along the cell boundary, local contractile stresses generated by class II myosins in the cortex may be transmitted to membrane proteins by their association with actin filaments in the cortex, generating actin-dependent nanoclusters(*14*, *28*, *29*). Therefore, we tested whether inhibiting class II myosin activity would influence the nanoclustering of a transmembrane protein with cytoplasmic actin-binding capacity. We used a chimeric transmembrane protein, FRTM-Ez-AFBD, where the extracellular domain of the Folate receptor (FR) coupled to the transmembrane region of the IgG receptor (TM) is linked to cytoplasmic actin filament-binding domain from Ezrin (Ez-AFBD) and has been shown to form actin-dependent nanoclusters(*14*). We employed Forster Resonance Energy Transfer between like fluorophores (homo-FRET), read out as changes in fluorescence emission anisotropy, to monitor the extent of nanoclustering of FRTM-Ez-AFBD(*30*) in cells expressing this protein (FRTM-Ez-AFBD cells). When labelled with a fluorescent analog of folate (Pteroyl-lysyl-BODIPY or PLB), nano-clustered proteins exhibit homo-FRET, resulting in a reduced fluorescence anisotropy compared to an actin-binding mutant of Ez-AFBD (R579A in FRTM-Ez-AFBD*) (*14*, *31*). Importantly, the reduction in anisotropy was also abrogated when class II myosin activity was inhibited with blebbistatin, a compound that interferes with its power stroke and reduces its ability to bind actin (*32*) (Fig. 2A,B). This is consistent with a loss in nanoclustering, as confirmed by the change in anisotropy obtained by photobleaching of the fluorophores. A reduction in anisotropy due to homo-FRET can be reversed by fluorophore dilution through photobleaching, characterized by a positive slope of the change in anisotropy as a function of photobleaching (see schematic in Supplementary Fig. S1A and ref. (*30*)). The positive slope observed with photobleaching PLB-labelled FRTM-Ez-AFBD signifies nanoclustering in contrast to the almost null slope obtained by photobleaching in blebbistatin treated cells, similar to that obtained for the actin-binding mutant protein, FRTM-Ez-AFBD* (Fig. 2C and Supplementary Fig. S1), confirming the loss of nanoclustering. Additionally, treatment of the cells with a cocktail of inhibitors, MLY (ML7+Y27632; against the regulatory light chain phosphorylating kinases ROCK and MLCK)(*33*), also resulted in a loss of FRTM-Ez-AFBD clustering (Supplementary Fig. S1E). Consistent with this data obtained with the synthetic actin-binding FRTM-Ez-AFBD, we have found that nanoscale patterning of various naturally occurring transmembrane proteins are also mediated by class II myosin activity(*28*, *29*). In particular, actin-dependent nanoclustering of cell-matrix adhesion protein CD44 and cell-cell adhesion protein E-Cadherin in *Drosophila* at the free cell-surface are perturbed by MLCK and class II myosin inactivation(*28*, *29*). Here the association of CD44 and E-Cadherin with actin is mediated by the cytosolic proteins, Ezrin and α-Catenin, respectively, that bind filamentous actin and the cytoplasmic tails of these proteins.

**Fig. 2.**
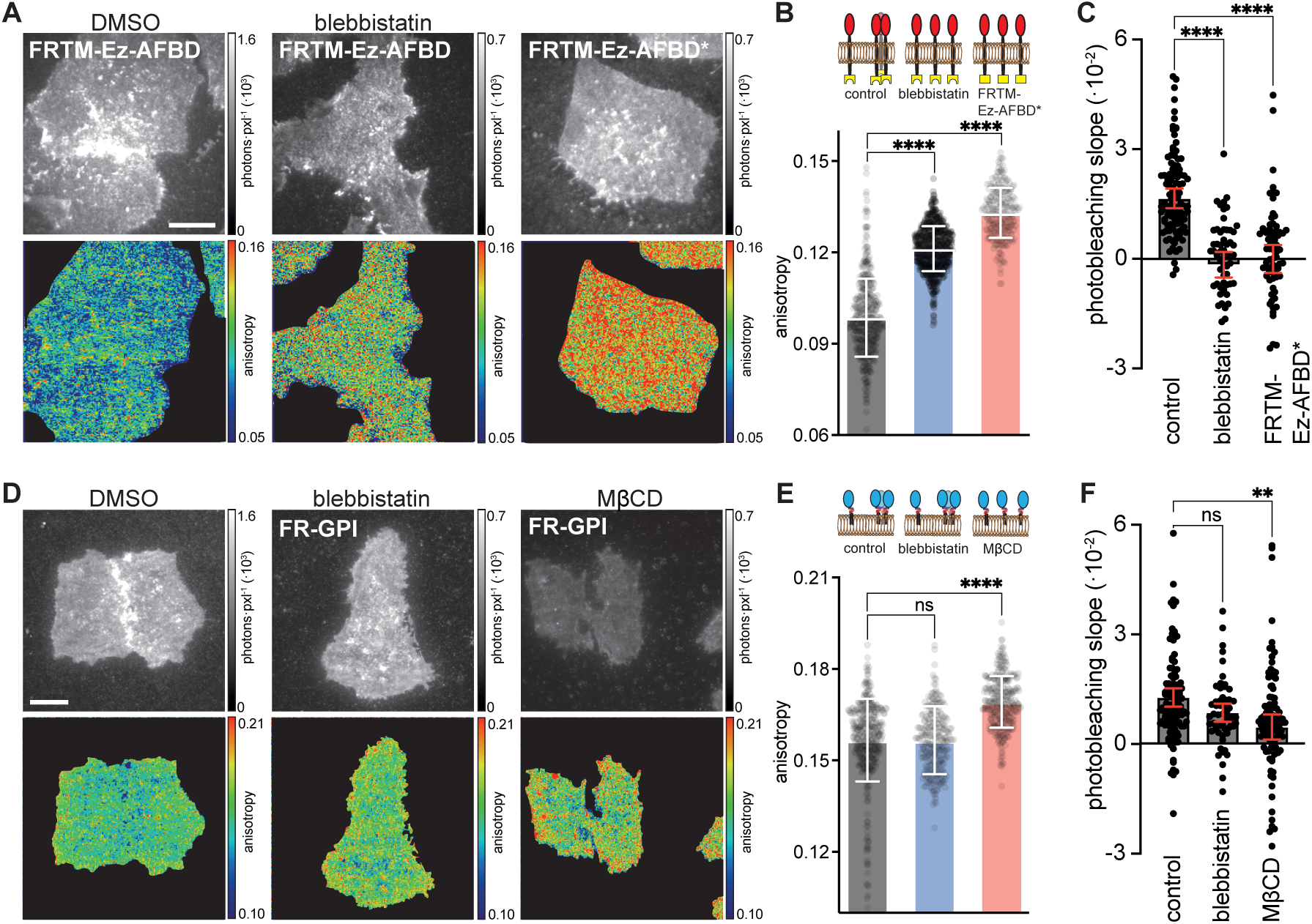
Effect of inhibition of class II myosin activity on membrane component organization. **A)** Intensity (top) and anisotropy (bottom) images were obtained using TIRF microscopy from cells expressing chimeric transmembrane proteins FRTM-Ez-AFBD or FRTM-Ez-AFBD*, labeled with PLB and treated with vehicle (DMSO) or blebbistatin (100 μM) as indicated. **B)** Bar graphs show the mean (±s.d) of emission anisotropy values (circles) collected from 4.5 μm^2^ regions of interest obtained from emission anisotropy images collected as in (a). Data for the control, blebbistatin and FRTM-Ez-AFBD* were obtained from 40 cells (446 ROIs), 51 cells (609 ROIs), and 13 cells (239 ROIs), respectively. **C)** Bar graphs show mean (±s.d) of photodilution slopes of the emission anisotropy from chimeric proteins following perturbations as in (a). Data for the control, blebbistatin and FRTM-Ez-AFBD* were obtained from 100 cells (N=7), 61 cells (N=4), and 70 cells (N=5), respectively. **D)** Intensity (top) and anisotropy (bottom) images obtained using TIRF microscopy from cells expressing FR-GPI and labelled with PLB and treated as indicated. **E)** Bar graphs show the mean (±s.d) of emission anisotropy values (circles) collected from 4.5 μm^2^ regions of interest obtained from emission anisotropy images collected as in (D). Data were obtained from 32 cells (396 ROIs), 19 cells (270 ROIs), and 25 cells (312 ROIs) for the control, blebbistatin and MβCD experiment, respectively. **F** Bar graphs show mean (±s.d) of photodilution slopes of emission anisotropy from FR-GPI following perturbations as in (E). Data were obtained from 109 cells (N=6), 55 cells (N=4), and 106 cells (N=6) for the control, blebbistatin and MβCD experiment, respectively. LUT, grey scale, total photons detected; color, anisotropy values. Scale bars, 10 μm. ****, **, and ns correspond to *p*-values of <10^-4^, <10^-2^, and >0.05, respectively.

We next addressed whether class II myosin activity also influenced the actin-driven organization of lipids in the membrane by examining the nanoclustering of GPI-APs, a representative of long-acyl-chain lipids at the outer leaflet (*19*, *31*, *34*). In stark contrast to the transmembrane actin-binding proteins, when cells expressing GPI-anchored folate receptor (FR-GPI) were treated with blebbistatin, there was minimal loss of clustering of the lipid-anchored protein (Fig. 2D-F). In *Drosophila* hemocytes, removal of class II myosin heavy chain (zip) or regulatory light chain (sqh) by UAS-GAL4 driven RNAi, also did not result in a loss of GFP-GPI clustering (Supplementary Fig S2A, B) whereas the actin-dependent clustering of the transmembrane *Drosophila* E-cadherin is lost(*29*). These results suggest that while nanoscale clustering of membrane proteins that are able to bind actin is mediated by class II myosins, outer-leaflet lipid-anchored molecules are likely to be clustered by trans-bilayer coupling and lateral stresses derived from different molecular agents.

### Class I myosins drive nanoclustering of GPI-anchored proteins

The presence of the lipid-binding evolutionarily conserved class I myosin isoforms juxtaposed to the cell membrane (Fig 1; Supplementary Fig. S2C, D) raises the possibility that these proteins may be involved in the formation of lipidic clusters that are not dependent on class II myosin function. There are eight human Myosin 1 isoforms (Myosins 1A, 1B,…,1H). Each isoform is a single heavy chain comprising of an actin-binding motor head domain, a calmodulin-binding neck region, and lipid-binding tail domain(*35*, *36*). We used the small molecule inhibitor, pentachloropseudilin (PClP) that decreases the interaction of class I myosin with cortical actin to inhibit class I myosin activity across all isoforms (*37*, *38*). When CHO cells expressing fluorescently labelled GPI-APs (FR-GPI or GFP-GPI cells), were treated with PClP there was an increase in the anisotropy of GFP-GPI (Fig 3A,B) and PLB-labelled FR-GPI (Supplementary Fig. S2E), compared to the vehicle treated control. Photobleaching of fluorophores confirmed that inhibition of class I myosin motor activity resulted in loss of homo-FRET, consistent with a loss of nano-clustering of GPI-APs (Fig 3C). By contrast, when FRTM-Ez-AFBD cells were treated with PClP, nanoclustering of the trans-membrane protein was unaffected (Fig 3D-F). These results were not cell-type specific; PClP-treatment also abrogated GPI-AP nanoclustering in U2OS and AGS cells (Supplementary Fig. S2E).

**Fig. 3.**
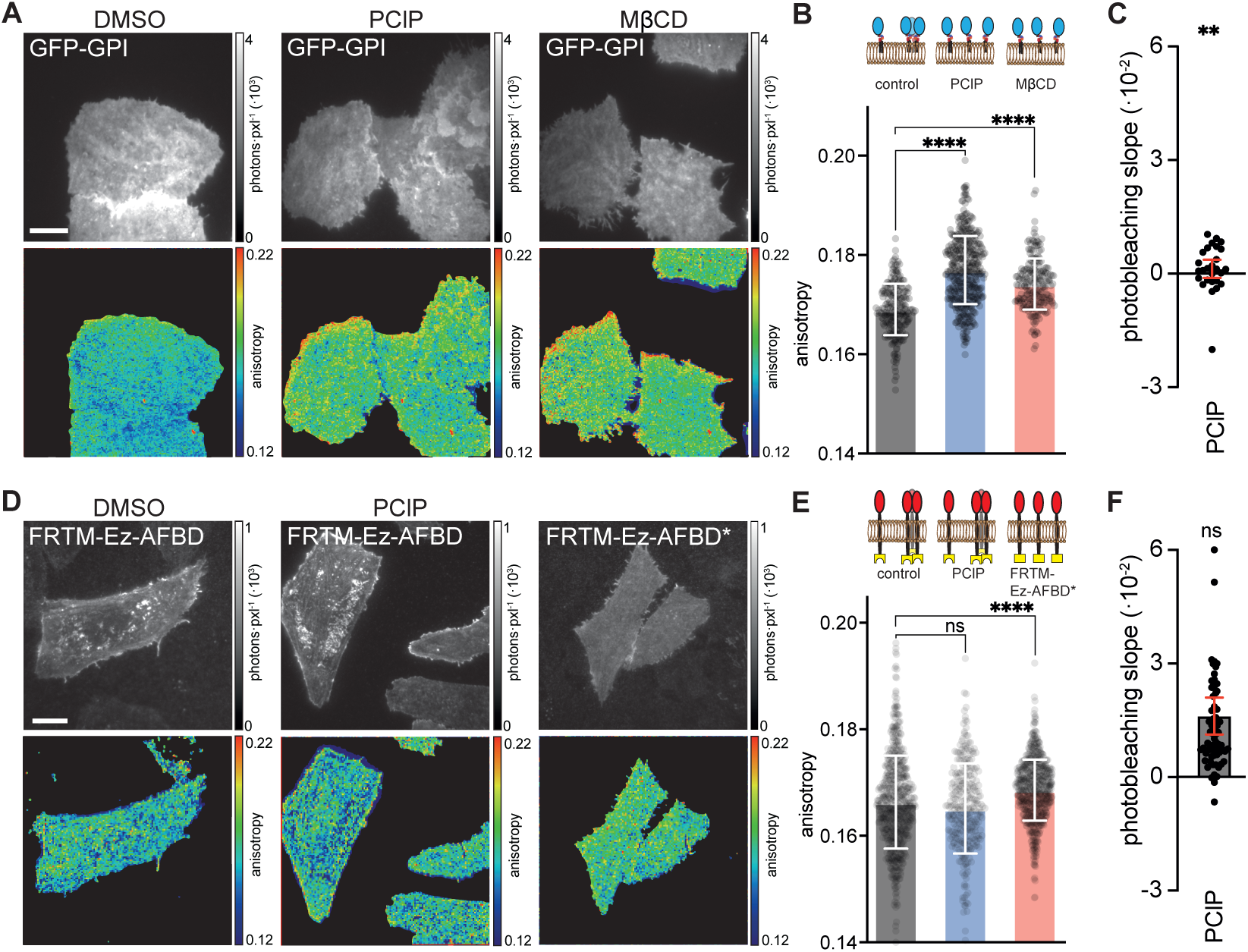
Effect of class I myosin inhibition on membrane component organization. **A)** Intensity (top) and anisotropy (bottom) images obtained using TIRF microscopy from cells expressing GFP-GPI, and treated with the vehicle (DMSO), myosin 1 inhibitor (PClP at 5 μM), or MβCD (10 mM). **B)** Bar graphs show the mean (±s.d) of emission anisotropy values (circles) collected from 4.5 μm^2^ regions of interest obtained from emission anisotropy images collected as in (a). Data were obtained from 48 cells (218 ROIs), 49 cells (415 ROIs), and 19 cells (205 ROIs) for the control, PClP and MβCD experiment, respectively. **C)**. Bar graph shows mean (±s.d) of the photodilution slopes of emission anisotropy from FR-GPI labeled with PLB, following PCIP treatment from 27 cells (N=3); *p*-value comparison is made with the control experiment in Fig 2f. **D)** Intensity (top) and anisotropy (bottom) images were obtained using TIRF microscopy from cells expressing chimeric transmembrane actin binding protein FRTM-Ez-AFBD or FRTM-Ez-AFBD* labelled with PLB, and FRTM-Ez-AFBD expressing cells were treated with the vehicle (DMSO) or PClP (5 μM). **E)** Bar graphs show the mean (±s.d) of emission anisotropy values (circles) collected as above from images as in (D). Data for the control, PClP, and FRTM-Ez-AFBD* were obtained from 50 cells (620 ROIs), 49 cells (299 ROIs), and 53 cells (683 ROIs), respectively. **F)** Bar graph shows mean (±s.d) of the photodilution slopes of emission anisotropy from PLB-labelled FRTM-Ez-AFBD from 68 cells (N=3) after treatment with PClP. *p*-value comparison is made with the control experiment in Fig 2c. LUT, grey scale, total photons detected; color, anisotropy values. Scale bars, 10 μm. ****, **, and ns correspond to *p*-values of <10^-4^, <10^-2^, and >0.05, respectively.

Class I myosin isoforms are expected to bind lipids via their tail domains and have a broad expression range across tissues, however, some isoforms such as MYO1E and MYO1F also bind proteins and others, MYO1A, MYO1G and MYO1H, are expressed only in specialized cell types and tissues(*35*, *36*, *39*). To identify which specific class I myosin isoform(s) may influence the general mechanism of GPI-AP clustering by connecting membrane lipids to the actin cytoskeleton, we examined the role of three of the lipid-binding, plasma membrane-localized isoforms that are widely expressed: MYO1B, MYO1C and MYO1D. In GFP-GPI-expressing U2OS cells, an increase in anisotropy of GFP-GPI was only observed if the levels of MYO1B, MYO1C and MYO1D were all simultaneously reduced by siRNA targeting all the isoforms (Supplementary Fig. S2F-J). This conclusively implicates class I myosins in lipid-based (GPI-AP) nanoclustering. This was further corroborated by UAS-GAL4 driven RNAi depletion of the orthologous Myo1C/H (Myo61F) and Myo1D/G (Myo31DF) in *Drosophila* hemocytes (Supplementary Fig. S2A,B).

### Class I myosins link cytoplasmic actin with outer-leaflet GPI-anchored proteins via PS

Class I myosins are associated with the inner leaflet via phosphatidylinositol bisphosphate (PIP2) and other negatively charged lipids such as PS via positively charged residues present in its tail(*40*). We have shown that lipids, including glycolipids and GPI-anchored proteins at the outer leaflet are coupled across the bilayer to PS via transbilayer interactions of long acyl-chain containing lipid species at both leaflets(*19*, *41*). The role of PS at the inner leaflet suggests that positively charged tail domain of class I myosins could provide an evolutionarily conserved molecular link (Supplementary Fig. S2C, D) between actin at the inner-leaflet and the outer-leaflet GPI-AP and other long-acyl chain containing glycolipids. The potential clustering of several PS lipids by the binding and accumulation of class I myosins may serve as the driving force for nanoclustering at the outer leaflet of the PM. To test this hypothesis, we created cell-attached blebs in FR-GPI-expressing cells and crosslinked the GPI-AP with antibodies(*19*). This generates large optically resolvable patches of GPI-APs(*42*) that can function as ’bait’ regions to recruit interacting ’prey’ proteins or lipids at the inner-leaflet (schematic in Fig 4A). GPI-AP crosslinking recruits PS lipids, as determined by co-localization of the PS-binding C2 domain of Lactadherin (LactC2-GFP; Fig 4B, C), consistent with previous studies(*19*). Significantly, class I myosins (GFP-Myo1B and GFP-Myo1C) were also recruited to the sites of cross-linked GPI-AP (Fig 4B, C). In contrast, when the transmembrane protein FRTM-Ez-AFBD* was cross-linked to serve as ‘bait’, GFP-Myo1C failed to localize to these patches (Fig 4B, C). These data reveal that class I myosins not only play a role in actively driving the clustering of GPI-APs, but they can function as coupling agents connecting cytoplasmic actin with PS at the inner-leaflet and GPI-APs and glycolipids at the outer leaflet, thereby behaving as ‘active linkers’.

**Fig. 4.**
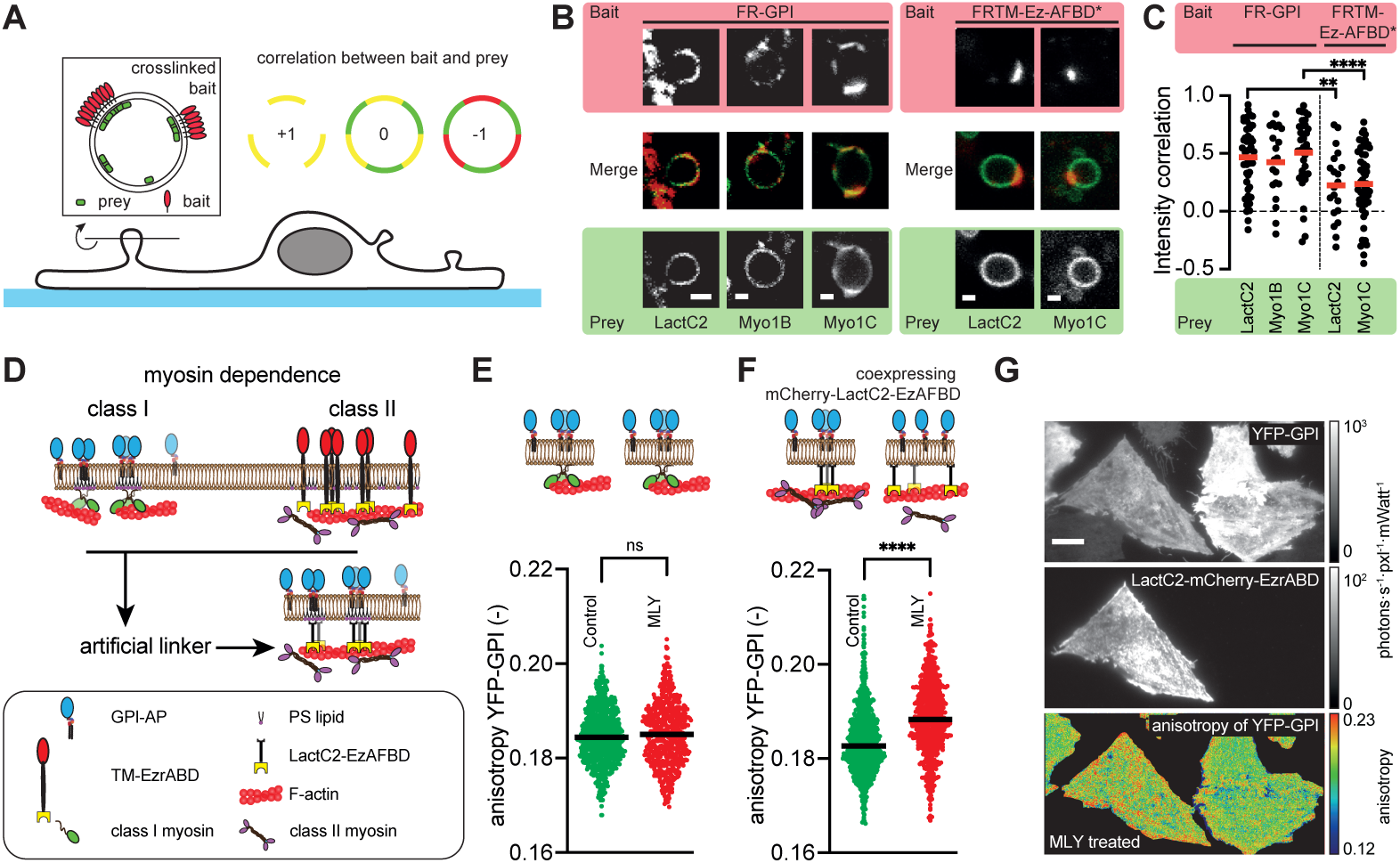
Class I myosins link outer-leaflet lipids via inner-leaflet PS to the cytoskeleton. **A)** Schematic representation of the assay to test the colocalization of outer-leaflet proteins (bait) and inner-leaflet (prey) proteins in cell-attached blebs. **B)** Representative confocal images of the equator plane of cell-attached blebs with either FR-GPI or FRTM-Ez-AFBD* as crosslinked bait and LactC2-GFP (PS-lipid binding protein), GFP-Myo1B or GFP-Myo1C as prey proteins at the inner leaflet. **c)** Gaphs shows quantification of the bait-prey intensity correlation index of the proteins visualized in **B** from multiple cell-attached blebs, where closed circles represent individual blebs and the red line corresponds to the mean value. **D)** Model for the bifurcation of the roles of class I myosin and class II myosin motors on association with membrane lipids and proteins, respectively generating distinct nanoscale clusters. Note that the synthetic construct containing both PS-binding and actin-binding would confer the ability of Myo2 to promote membrane lipids reorganization. **E,F)** Graphs showing the differential sensitivity of YFP-GPI clustering to a myosin class II inhibitor cocktail, MLY (20 μM ML7 and Y27632) in the absence or presence of the synthetic PS- and actin-binding linker, LacC2-EzAFBD. Data were obtained from 77 cells (1081 ROIs) and 105 cells (1041 ROIs) in (e), and 54 cells (1108 ROIs) and 53 cells (649 ROIs) in (f), for the control and MLY perturbations, respectively. **G) Intensity (top and middle) and anisotropy (bottom) images of YFP-GPI cells, co-expressing mCherry-LacC2-EzAFBD (middle) exhibiting sensitivity to the MLY cocktail.** Scale bars, 2 μm (B) and 10 μm (G). ****, **, and ns correspond to p-values of <10^-4^, <10^-2^, and >0.05, respectively.

We next asked if we could redirect GPI-APs to interact with the class II myosin-driven clustering mechanism by linking PS to an actin-binding domain. To this end, we used a synthetic PS-actin linker consisting of LactC2 and the Ez-AFBD (LactC2-EzAFBD) (see schematic Fig 4D). CHO cells expressing YFP-GPI were transiently transfected with mCherry-LactC2-EzAFBD, and the emission anisotropy of YFP-GPI in transfected and un-transfected cells were compared for their sensitivity to class II myosin activity by blebbistatin treatment (Fig 4E, F and Supplementary Fig. S3). Consistent with a redirection of the clustering mechanism we observed that the anisotropy of YFP-GPI, developed sensitivity to inhibition of class II myosin activity only in cells co-expressing LactC2-EzAFBD (Fig 4E-G).

### Nanocluster-rich domains of GPI-anchored proteins and transmembrane actin-binding proteins are spatially segregated

Our results thus far provide evidence for at least two types of active clustering pathways in the membrane; one localized at the membrane where class I myosin drives the nanoclustering of lipid-linked membrane components, and another mediated by the more cortically localized class II myosins that generate nanoscale clusters of transmembrane proteins linked to actin. Our earlier work showed how nanoscale clustering of membrane components was a natural consequence of the contractile stresses applied by actomyosin(*14*). Since the molecular agencies for these contractile stresses are in different strata of the cortex we asked if there is a segregation at the nanoscale (see schematic Fig 5A). To address the issue of co-clustering or segregation at the nanoscale we co-expressed both FR and GFP-linked to TM-Ez-AFBD or GPI anchors. In cells expressing both FRTM-Ez-AFBD and GFP-TM-Ez-AFBD, the anisotropy value of FRTM-Ez-AFBD increased as a function of the expression level of GFP-TM-Ez-AFBD (Fig 5A; red data and line in graph). This indicates that GFP-TM-Ez-AFBD stochastically ‘diluted’ FRTM-Ez-AFBD in the nanoclusters. Similarly, when FR-GPI and GFP-GPI were co-expressed, the extent of homo-FRET of either of the two was inversely proportional to the relative expression level of the other, consistent with co-clustering of the two proteins at the nanoscale (*31*). By contrast, there is no co-clustering when FRTM-Ez-AFBD and GFP-GPI are co-expressed (Fig 5A; black data and line in graph). This shows that at the cell surface the diverse myosin machineries build distinct nanoscale clusters.

**Fig. 5.**
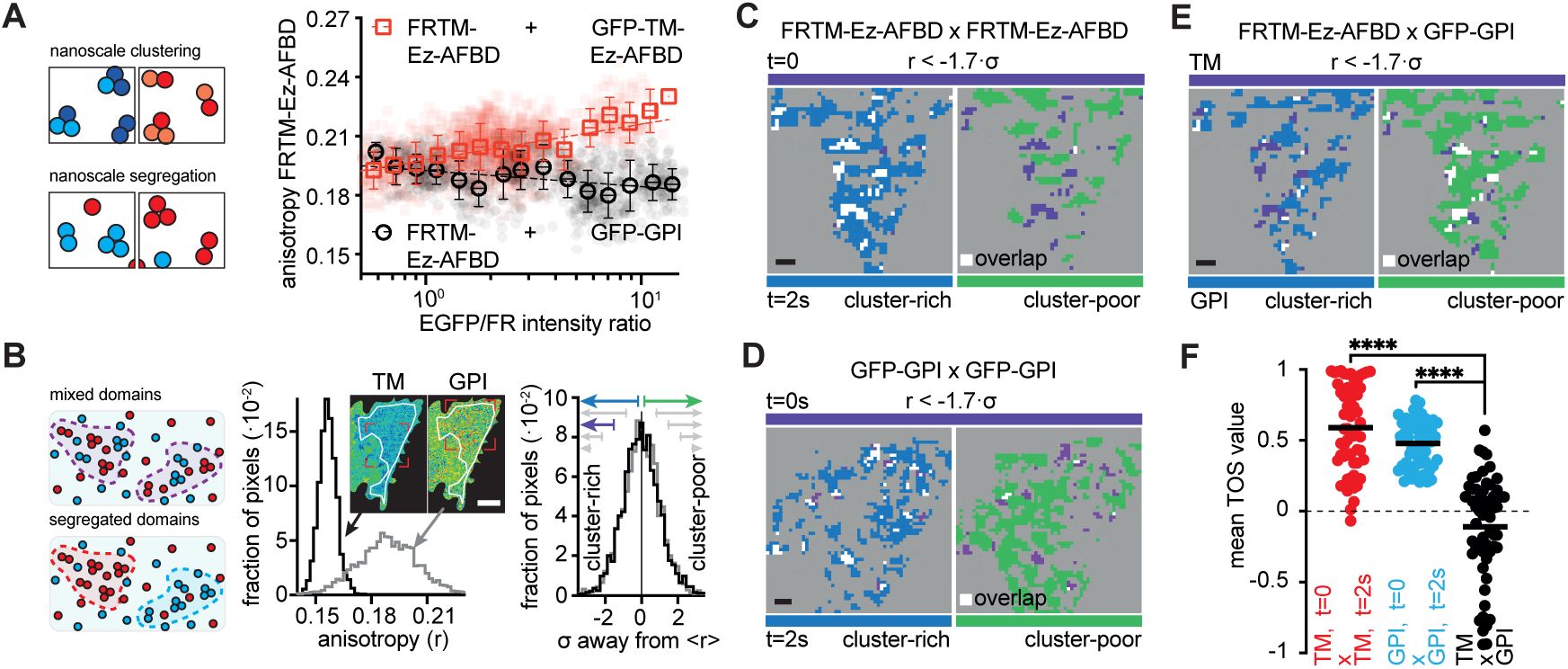
Spatial sorting of myosin activity zones. **A)** Schematic (left) of nanoscale (co)clustering, and plot (right) showing the change in anisotropy as a function of the ratio of GFP to PLB-labelled FRTM-Ez-AFBD in cells co-expressing both proteins. Note that anisotropy increases when GFP-TM-Ez-AFBD is co-expressed with FRTM-Ez-AFBD, red (1421 ROIs from 67 cells, N=2), square data points were obtained by averaging emission anisotropy of FRTM-Ez-AFBD at the indicated GFP/FR ratio bin (±s.d.). However, there is no change when GFP-GPI is co-expressed with FRTM-Ez-AFBD, black (1021 ROIs from 59 cells, N=2), open circles are mean±s.d. of the anisotropy at the different ratio bins. **B)** Schematic of mesoscale domain organization (left) and histograms of the raw (middle) and normalized (right) pixel anisotropy values of the same selected cell (see anisotropy image insets) for FRTM-Ez-AFBD (black line) and the GFP-GPI (gray line) signals. **C-E)** Threshold images of pixels corresponding to cluster-abundant pixels (< -1.7·σ, purple pseudo colored) in one image together with the cluster-rich (< 〈*r*〉, blue pseudo colored) or cluster-poor pixels (> 〈*r*〉, green pseudo colored) of the corresponding second image: temporal FRTM-Ez-AFBD correlation (C), temporal GFP-GPI correlation (D), and FRTM-Ez-AFBD with GFP-GPI crosscorrelation (E). Note the abundance of overlapping pixels (white) with cluster-rich regions and their lack with cluster-poor regions for the temporal correlations of the same component as compared to the inverse trend for the crosscorrelation. **F)** Graph of the mean TOS-value between FRTM-Ez-AFBD and GFP-GPI. Their averaged temporal and crosscorrelation TOS-values have been obtained from the area enclosed by the dashed line in the TOS matrix of Supplementary Fig. S5E). LUT, color, anisotropy values. Scale bars, 10 μm (images in B) and 2 μm (pixels maps in C-E). **** corresponds to a p-value of <10^-4^.

At the mesoscale we visualized the distribution of active emulsions enriched in TM-Ez-AFBD and GPI-AP nanoclusters, using near-simultaneous fluorescence emission anisotropy maps obtained from PLB-labeled FRTM-Ez-AFBD and GFP-GPI expressing cells. Nanocluster-rich active emulsions (low anisotropy regions) appear as domains spanning several pixels with a characteristic scale of about ∼450 nm(*14*, *16*). To study the relationship between the different types of active emulsions (Fig. 5B) we established an analytical pipeline to explore correlations between these domains. This utilizes a method that calculates threshold overlap scores (TOS)(*43*) to quantitatively assess the extent of non-random overlap between two different distributions (Fig 5B, see also Supplementary Information and Supplementary Fig. S4). In living cells, using such a metric there appears to be a significant overlap of cluster-rich mesoscale domains of FRTM-Ez-AFBD or GFP-GPI in images derived from sequential frames spaced 2 seconds apart (Fig 5C, D, Supplementary Fig. S4C-E). When analyzed across many cells and fields, this is reflected as high mean TOS values (TOSFRTM-EzAFBD=0.59±0.31 and TOSGFP-GPI=0.48±0.16; Fig 5F and Supplementary Fig. S4E). This indicates a stable network of such domains formed by the two different myosin motors. However, cluster-rich domains of GFP-GPI exhibited a small but significantly negative overlap score (TOS=-0.11±0.37) when correlated to images of the cluster-rich domains of FRTM-Ez-AFBD derived from sequential frames in the same cells (Fig 5E and F, and Supplementary Fig. S4E). This is consistent with the segregation of these mesoscale regions built by the different myosin motors in space as depicted in schematic in Fig. 5B.

### Active Flory-Huggins theory predicts segregated domains generated by active stresses with tunable concatenation

To understand the basis for the mesoscale patterning of membrane components driven by multiple membrane localized active stresses we elaborated an active Flory-Huggins (AFH) theory, originally built to describe these active emulsions built by a single contractile agent(*26*). Here we extend this analysis to incorporate two agents of contractility, namely class I and II myosin that drive the clustering of class I myosin-binding *lo* components and the class II myosin-associated actin binding component, FRTM-Ez-AFBD, respectively. For simplicity, we formulate the AFH theory in terms of the dynamics of the area fractions of the *lo*-preferring component (GPI-AP) and TM-Ez-AFBD on the membrane in juxtaposition with the actomyosin cortex. The dynamics is driven by forces that arise from (i) passive interactions between the components and (ii) active stresses on the *lo* component and TM-Ez-AFBD arising from class I and II myosin, respectively.

In turn, active stresses depend on the local concentrations and contractile flows of class I and II myosins in the cortex as described by active hydrodynamics equations(*26*), that incorporate their differential turnover, competitive binding and contractile activity (see schematic Fig. 6A; Supplementary Information and Supplementary Fig. S5). The theory predicts that the *lo* component segregates from the bulk *ld* phase even at T> Tc, as verified in earlier experiments(*16*), driven by contractile flows associated with the myosin activity linked to the *lo* component. In addition, contractile flows associated with class II myosin, drive TM-Ez-AFBD into forming distinct domains (Fig. 6B and Supplementary Fig. S5). Mesoscale domains of each component, defined for the numerical simulations as regions with area fractions above a threshold (0.3) show minimal overlap where the *lo* component and TM-Ez-AFBD do not interact (Fig 6B), consistent with experiments (Fig. 5B-F). Interestingly, unlike usual phase segregation, these mesoscale domains are not homogeneous but show granularity, with smaller regions of high density, identified with nanoclusters. The theory predicts that the extent of overlap (concatenation) at the mesoscale depends on competitive binding and contractile activity of the myosins, through the parameter Q (Fig. 6C, right panel, see also Supplementary Information). Moreover, the theory predicts that the extent of concatenation of the distinct mesoscale regions can be enhanced by increasing the in-plane affinity between their components (Fig. 6B, 6C left panel, and Supplementary Fig. S5).

**Fig. 6.**
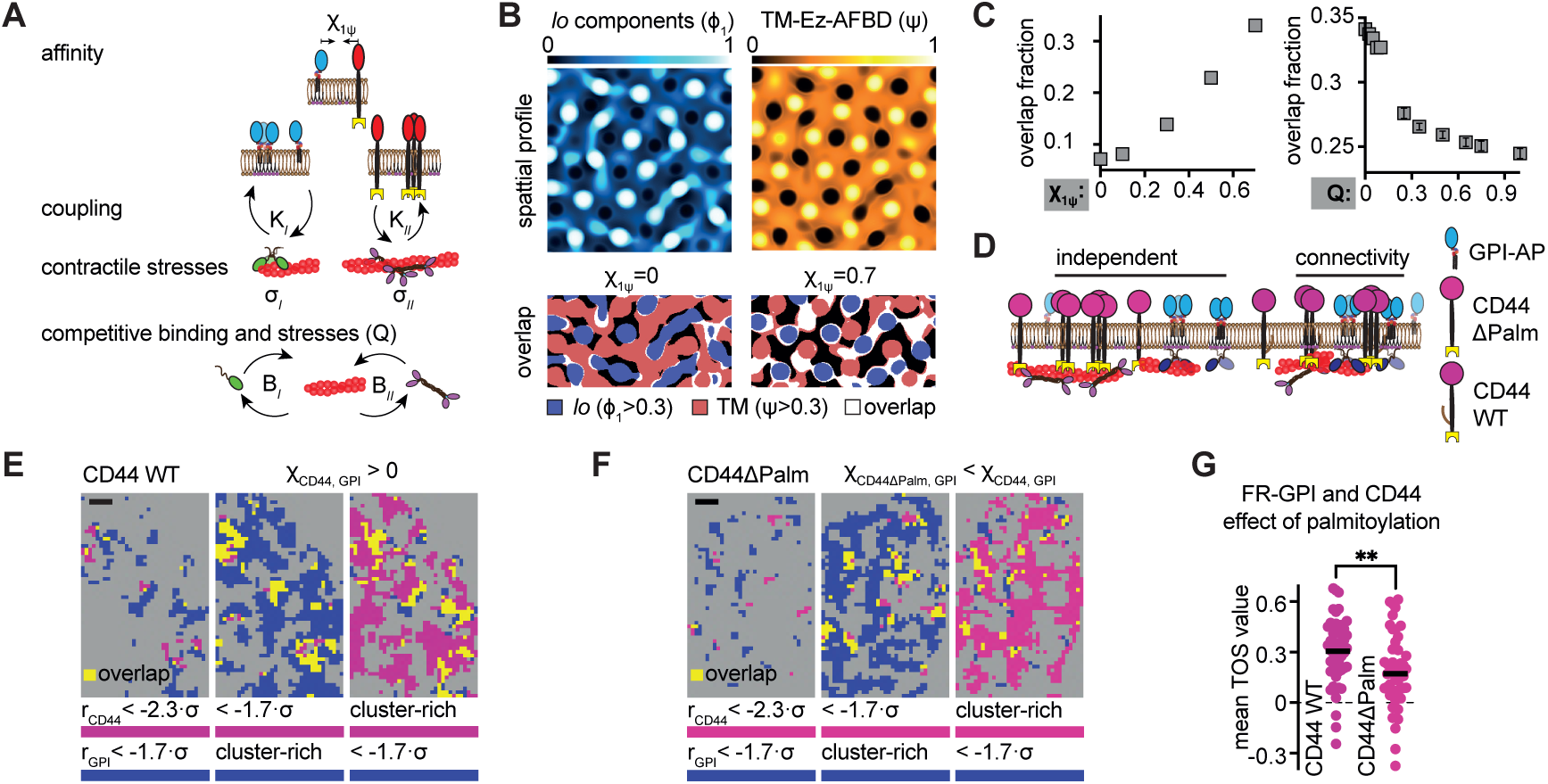
Active Flory-Huggins theory and concatenation of myosin activity zones. **A)** Elements of the components for the active Flory-Huggins (AFH) theory incorporating two agents of contractile stresses (σ*_I,II_*). The theory can be separated into (1) an acto-myosin component where competitive myosin binding and stresses determine the local accumulation of active class I or class II components and (2) in-plane interactions strength (χ_1ψ_) between the different membrane components coupled to distinct stress agents (See also Supplementary Information). **B)** Representative area-fraction images resulting from numerical simulations of the AFH theory for the *lo*-phase (top left) and TM-Ez-AFBD (top right). Bottom row: two representative overlap images for mesoscale domains (area fraction >0.3) where the interaction parameter (χ_1ψ_)) between the *lo*-phase and TM-Ez-AFBD were set at 0 (left) and 0.7 (right). **C)** Fraction overlap as a function of interaction parameter (left) or competitive stresses between the two classes of myosin (right). **D)** Cartoon displaying the possibility for connectivity leading to concatenation of mesoscale domains organized by the different classes of myosin motors. **E-F)** Zoomed-in thresholded images of pixels corresponding to cluster-abundant pixels (CD44: -2.3·σ or -1.7·σ, purple pseudo colored; GPI:-1.7·σ, blue pseudo colored) and cluster-rich pixels (CD44: < 〈*r*〉, purple pseudo colored; GPI: < 〈*r*〉, blue pseudo colored). **G)** Graph of the mean TOS-values between CD44 and FR-GPI with and without the palmitoylation sites. LUT, color, anisotropy values. Scale bars, 2 μm (pixels maps in E-F). ** corresponds to a p-value of <10^-2^.

To test this prediction we determined if the distinct types of active emulsions could become concatenated(*44*) by increasing the mutual chemical affinities between the components of the two types of active emulsions (schematic in Fig. 6A, D). Palmitoylation is a common reversible post-translational modification of transmembrane proteins which increases the affinity of the transmembrane species for *lo* domains (*45–48*). Consistent with our prediction we find an enhanced spatial overlap of palmitoylated CD44-GFP emulsions with the GPI-AP emulsions (TOS=0.30±0.19; Fig. 6E, G); mutation of the palmitoylation sites in CD44 (CD44!′1Palm-GFP) led to a loss of this spatial correlation (TOS=0.17±0.25; Fig. 6F, G). Taken together, these data imply that the bifurcation of non-equilibrium membrane organization by active stresses arising from the two classes of non-muscle myosins, provide a chemically reversible mechanism to switch between segregated and concatenated states of mesoscale domains (see model in Fig. 7).

**Fig. 7.**
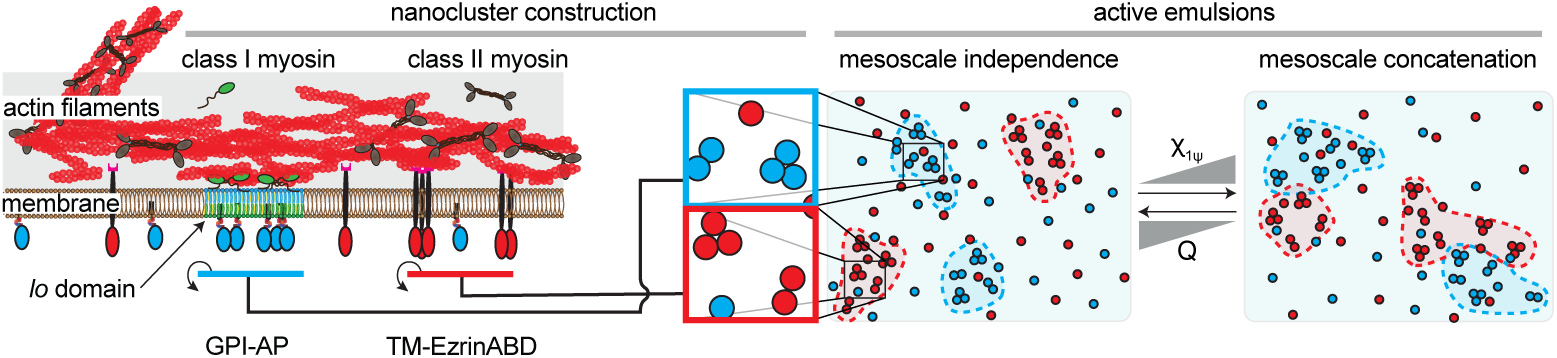
Model. Schematic showing the stratification of myosin motors at the plasma membrane driving the formation of nanoscale clusters of distinct components. These nanoclusters assemble into segregated mesoscale active emulsions that may be reversibly concatenated by increasing in-plane interactions strength (χ_1ψ_) between the different membrane components even if they are coupled to different stress agents.

### Functional implications of actively segregated nanoclusters

To address a functional consequence for the mechanistic bifurcation of the two myosin families in building domains enriched in the distinct types of nanoclusters, we turned to cell-spreading in fibroblasts. This process provides an ideal functional context for testing the roles of the distinct myosin requirements because membrane recruitment of class I myosins and activation of class II myosins are temporally separated during cell spreading(*49*, *50*) (see schematic Supplementary Fig. S6A). Crucially, cell-spreading is contingent on the generation of active emulsions enriched in GPI-AP nanoclusters right at the onset of spreading (*51*) preceding the timing of myosin 2 activation (Supplementary Fig. S6A). From the cell spreading area montages (Fig 8A and Supplementary Fig. S6B) it is evident that inhibition of class I myosin also abrogates the cell spreading response. A closer look at the cell spreading response (Fig. 8B) displays both an increase in the lag-time before cell spreading (Fig. 8C), and a decrease in the extent of area increase during spreading (Fig. 8D), consistent with class I myosin activity being required for the generation of GPI-AP emulsions downstream of activation of the integrin receptors. By contrast, inhibition of class II myosin does not increase the lag-time nor abrogate cell spreading responses (Fig. 8A-D); it augments the extent of spread area, likely by reducing cortical contractility(*52*). These results highlight the role of the two classes of non-muscle myosins in functional segregation of specific components of the cell membrane.

**Fig. 8.**
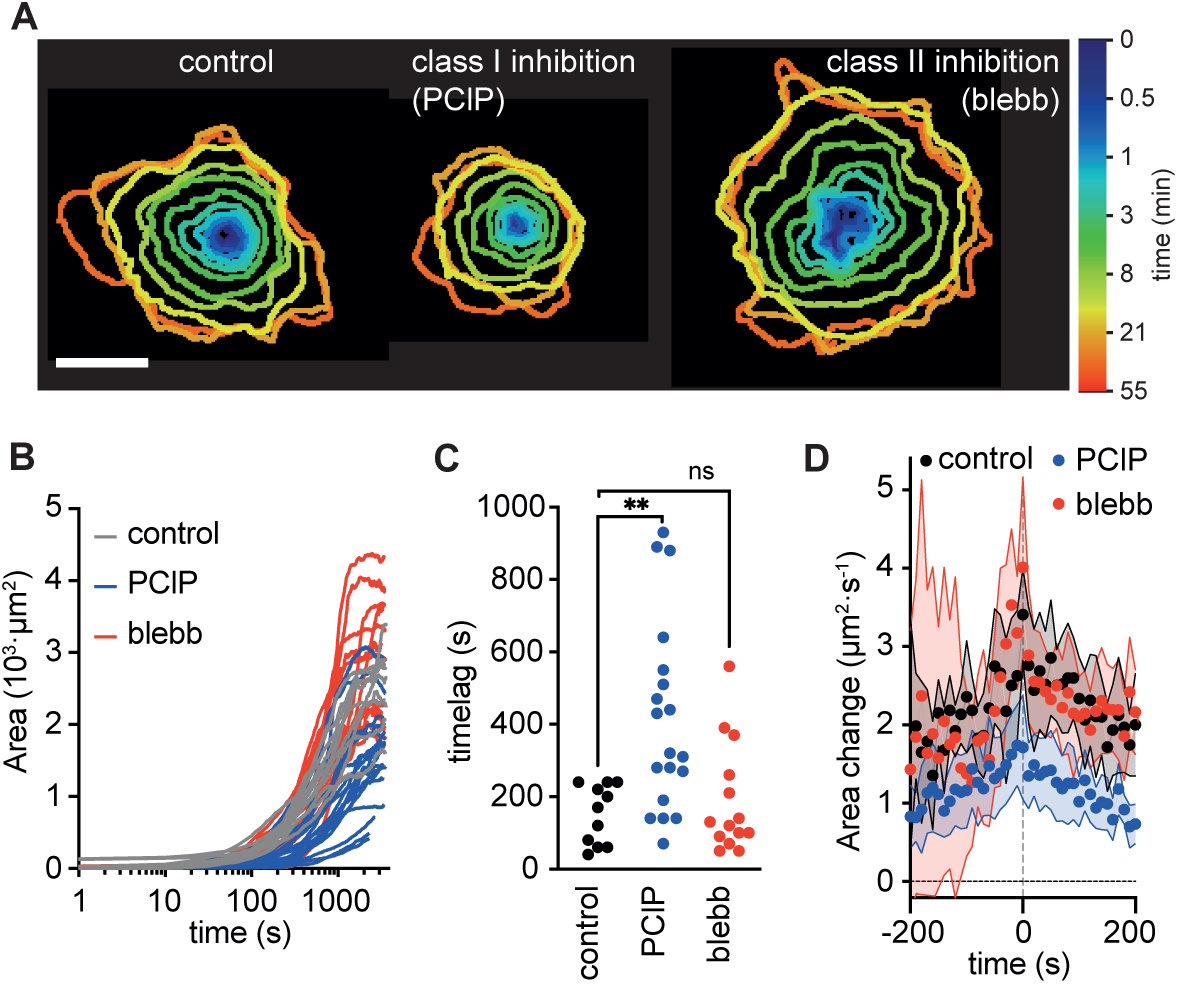
Functional consequences of myosin class independence membrane patterning on cell spreading. **A)** Temporal evolution of spread area from cells treated with vehicle (DMSO, 1 hour), PCIP (5 μM, 1 hour) or Blebbistatin (blebb 50 μM, 1 hour) and adhering to fibronectin-coated surfaces imaged using Interference Reflection Microscopy. **B-D)** Graphs showing the time-dependent cell spread area per cell (B), time lag between cell attachment and cell spreading (C) and the extent of cell area increase (D; mean ± s.d) during spreading of the same cells as in B where the maximum rate of area increase was shifted to t=0. These data have been obtained from 11, 14 and 19 cells for DMSO, blebb and PClP, respectively, N=3. Scale bar, 10 μm. **, and ns correspond to *p*-values of <10^-2^, and >0.05, respectively.

## Discussion

Nanoclusters such as those described here are emerging as major organizational units of larger scale membrane platforms for signal transduction(*53*, *54*), cell-cell interactions(*29*, *55*), cell migration(*56*), and many other cellular processes(*57*, *58*). We provide evidence that multiple cell membrane constituents are patterned by distinct classes of ancestral myosin motors at the nano and mesoscales. A stratified actomyosin cortex appears as a strategy for functional and tunable segregation of components in the membrane. Class II myosins, embedded in the actin cortex transmit contractile stresses at a distance to membrane proteins attached to dynamic actin filaments in the cortical actin meshwork generating nanoclusters and emulsions of the same. On the other hand, class I myosins when directly coupled to the membrane with their ability to recruit and move actin, generate lipidic nanoclusters leading to functionally active emulsions at the mesoscale. Identifying such motorized plasma membrane-cytoskeleton linkers solves a long standing question in the field, of how outer leaflet lipids and proteins sense and register underlying actomyosin contractility to cluster at the plasma membrane. Given that class I myosin associates with PS at the inner leaflet and couples via a trans-bilayer interaction with outer leaflet GPI-APs to create *lo-* ordered nanodomains(*59*, *60*), it is not suprising to see the generation of mesoscale *lo-*domains. This exemplifies the capacity of such an active process in building membrane regions of distinct chemical composition and properties.

This capacity emerges from a common physical mechanism connected to active stresses of polar filaments derived from their engagement with diverse myosin motors. The fact that several ATP consuming machines (actin and at least two myosin classes) pattern membrane components into nanoscale clusters and consequently build active emulsions, foregrounds the role of non-equilibrium mechanisms in structuring and organizing the PM. The mechanism described here (Figs. 6A and 7) also provides a tunable way to concatenate these domains via reversible post-translational modifications (*e.g.* palmitoylation) that confer weak affinity interactions between components of the distinct emulsions(*44*).

The patterning potential of the mechanism presented here is far-reaching; the association of class I myosins with specific lipids (*i.e.* PS and PIP2 for Myosin 1C) and membrane receptors (*i.e.* Myosin 1B to EphRB(*61*)) and the class II myosin-infused cortex with many proteins with actin-binding capacity, are just some examples. A major contribution to this canvas is also the broad diversification potential of the proposed active mechanisms at the level of actin binding, positioning and mechanics(*62*). We speculate that the segregation potential of this ATP-fueled fabric is made available for the crucial purpose of modulating information transduction, because it can not only be regulated in space and time but can also enable chemically reversible concatenation of diverse domains during the construction of signaling cascades(*58*).

## Materials and Methods

### Materials

Tables listing reagents and their source, cell lines, pDNAs, *Drosophila* strains, and simulation parameter are provided in the Supplementary Tables S1-S5.

### Methods

#### Cell culture, transfection and labeling for live imaging

Transmembrane or GPI-anchored fluorescent reporters were stably expressed in Chinese hamster ovary (CHO) cells(*14*, *31*, *34*). CHO-derived cell lines were maintained in Ham’s F12 media supplemented with 10% FBS, 1% penicillin-streptomycin-glutamine cocktail and with appropriate selection antibiotics. The maintenance media was free of phenol red and where the folate receptor domain was used for labeling the maintenance media was additionally folate free. The FBS used for these cells was dialyzed as described earlier(*34*). Human U2OS cell lines, U2OSGG stably expressing GFP-GPI(*51*), were maintained in McCoys, supplemented with 10% FBS, 1% penicillin-streptomycin-glutamine cocktail and with appropriate selection antibiotics.

Cells were plated in 35 mm glass-bottom dishes, 48 hours prior to live imaging, in maintenance media without selection antibiotic. Where applicable, transient transgene expressions were carried out by transfection of cells with Fugene6 following manufacturer’s guidelines. When required, human Fibronectin (hFN) coated dishes were prepared by incubating the glass bottom dish for 1 hour at 37°C with 10 µg/ml hFN in PBS.

Cells were imaged in M1 buffer (20 mM HEPES, 150 mM NaCl, 5 mM KCl, 1 mM CaCl2 and 1 mM MgCl2, adjusted to pH 7.4)) supplemented with 2 mg/ml glucose (M1G). In case of fluorescent protein tagged membrane proteins, cells were incubated with 100 µg/ml of cycloheximide in culture media for at least 3 hours prior to imaging. For drug and inhibitor treatments the cells were incubated at the desired concentration and indicated time before bringing the dish to the microscope for imaging. Folate receptors were labeled by incubating the cells with 400 nM pteroyl-lysine-BODIPY-TMR (PLB) in M1G at 37°C degrees for 10 minutes as the final step in presence of the inhibitor or its absence.

#### Measuring relative localization of class I and class II myosins

CHO cells that were plated for 2 days, were co-transfected and grown for an additional 12-16 hours with pDNAs encoding GFP-Myo1C and mCherry-Myo2A, de-adhered with TrypLE, pelleted down and replated on hFN-coated dishes. Replating ensured that most cells were individually adhered and did not have cell-cell contact regions, which allowed optimal optical segmentation of single plasma membrane slices. After 2 hours the cells were labeled with MemBrite Fix 405/430 following the manufacturer guidelines and subsequently fixed with 2% PFA in M1G for 20 min at room temperature. After a permeabilization step with 0.1% TritonX-100 for 15 min the cells were labeled with 200 nM Abberior STAR-635-phalloidin, washed three times and imaged on a Zeiss LSM880 confocal microscope equipped with an Airyscan detector using a 63X 1.4NA oil objective. Because only two dyes can be detected during a single Airyscan z-stack, each cell was imaged using 3 spectral combinations (with Zeiss Airyscan deconvolution lateral and axial pixel sizes): GFP-UV (35.3 nm and 140 nm), GFP-Far Red (42.5 nm and 170 nm), Far Red-mCherry (48.9 nm and 180 nm). Using 100 nm TetraSpeck beads the three combinations were corrected for any spatial offsets. Next, between 2-9 line profiles of around 6 µm in length were selected per cell going from extracellular to the cell interior passing the plasma membrane. For each line profile the peak, corresponding to the position of the labeled component, was determined by fitting with a Gaussian function and the lateral shift between the components was calculated. Starting from the membrane label (UV) the Myo1c position (GFP) was determined (*d_mem→myo1c_*), next the phalloidin labeled F-actin position (Far Red) was determined with respect to Myo1c (*d_myo1c→f−actin_*), lastly the Myo2a (mCherry) position with respect to the F-actin was measured (*d_f−actin→myo2a_*). All the measured distances were calculated with respect to the membrane position (Fig. 1d). Each extracted line profile was shifted with respect to the membrane peak position at 1 µm intervals and all 69 line-profiles were averaged and their mean ± standard deviation displayed in Fig. 1c.

#### Inhibitor treatments

For cholesterol depletion, cells were exposed to 10 mM MβCD in M1G for 45 min with three buffer changes every 15 min at 37°C. Myosin class II was inhibited using either 100 µM Blebbistatin or a cocktail that includes 20 µM of ML7 and 20 µM Y-27632 in M1G for 1 hour at 37°C. Class I myosins were inhibited by incubating the cells for 1 hour with 5 µM of PCIP(*38*, *63*) at 37°C.

#### Steady-state emission anisotropy measurements

Emission anisotropy experiments were performed on a spinning disk confocal microscope or on a total internal reflection fluorescence (TIRF) microscope as detailed previously(*30*, *64*). Emission anisotropy images were collected by splitting the parallel (*I*_∥_) and perpendicular (*I*_⊥_) emission using a polarizing beamspliter and collecting the data on two cameras simultaneously.

The values at each pixel in the images that were acquired were all converted to photo-electrons. For data recorded with EMCCD cameras the experimental EMgain-factor was calculated directly from the image using the method of Heintzmann et al.(*65*) The pixel-wise gain for the sCMOS was obtained using an experimentally pre-calibrated gain map(*66*). The raw image signal (*I*_∥_ and *I*_⊥*system biased*_) was subsequently calculated by subtracting the experimentally determined dark image and dividing the image by the gain-value (EMCCD) or by the gain-image (sCMOS). The raw images from the two polarisations were spatially aligned using iterative image registration combined with an affine transformation function (Matlab). Alignment corrections were kept at a minimal by experimentally aligning the images of sub-resolution beads before each imaging session. To correct for any system dependent bias in the polarisation detection due to the optics, a *G_factor_* image was determined, using a 1 μM Rho6G solution where *G_factor_* = *I*_∥,*Rho*6*G*_ / *I*_⊥,*Rho*6*G*_: and the raw signal at the perpendicular channel (*I*_⊥*system biased*_) needs to be corrected for this bias: *I*_⊥_ = *G_factor_*· *I*_⊥*system biased*_. After these corrections the anisotropy was determined following:

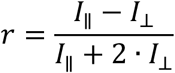

To reduce the error in the anisotropy representation both the *I*_∥_ and *I*_⊥_ were binned by summing up three by three neighbouring pixels, improving the signal 9-fold. The calculated anisotropy image was masked, showing only anisotropy values at pixels where there was signal above the background. The emission anisotropy values were obtained from the summed number of *I*_∥_ and *I*_⊥_ photons of square regions-of-interest (ROI) of 2.1 μm vertices length. Intensity flat regions from the plasma membrane were selected randomly, only avoiding regions close (<2 μm) to the cell edge. The corrected signal inside of these ROIs was summed for either *I*_∥_ and *I*_⊥_ and the anisotropy (*r*) and total intensity (*I_T_* = *I*_∥_ + 2 · *I*_⊥_) calculated. Plotting *r* versus *I_T_* allowed for the determination of the intensity region that is free from noise and trivial (concentration dependent) FRET (*30*). For each experiment, comparable regions were selected and displayed as a bar graph together with all ROI values and comparison was done using the Mann-Witney ranked test (Fig. 2B,E, Fig. 3B,E, Fig. 4E,F and Supplementary Fig. S2D-F,I).

#### Emission anisotropy upon photobleaching

A powerful method to determine whether the measured depolarization of the emission anisotropy is a result of the occurrence of h-FRET is to follow the anisotropy during photobleaching. During the photobleaching process, fluorophores undergo photobleaching in a stochastic manner, therefore the number of unbleached fluorophores in the same nanocluster reduces (Supplementary Fig S1A). For small cluster-sizes as in the case of the nanoclusters under study, the measured anisotropy therefore increases linearly with the photobleaching extent(*30*, *31*); for the situation where there are only monomers, there is no change in the emission anisotropy. Only fluorophores that cannot function as an energy acceptor after photobleaching can be used for these experiments, thus this excludes extracellular positioned GFP because in non-reducing environments the chromophore gets trapped in a dark-state that can still participate in energy migration process(*67*). Therefore, all our photobleaching experiments were performed with the PLB-labeled folate-analogues (see example sequences in Supplementary Fig S1B,C) where the fluorophores were photobleached using 561 nm laser power at 20-30 mWatt and 50-100 frames (100-200 ms integration time) were acquired at 0.5-2 Hz to measure emission anisotropy during this process. The intensity trace from each ROI, chosen from regions that were relatively uniform in intensity and had minimal amounts of bright spots, was background corrected and the calculated anisotropy was plotted with respect to the photobleaching extent (Supplementary Fig S1D), defined as 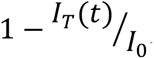, with *I_T_*(*t*) as the total intensity at frame t and *I*_<_ being the total intensity at frame t=0. A line was fitted to each ROI time-series with “*Photobleaching extent*” <0.6 and the slopes were recorded (Supplementary Fig S1D, E). Data on the slope per cell were combined from multiple cells and experiments and compared using the Mann-Wittney rank test (Fig 2C, F; Fig 3C, F and Supplementary Fig S1E).

#### *Drosophila melanogaster* RNAi knock down and GFP-GPI anisotropy

*Drosophila melanogaster* flies were maintained at 22 °C in glass vials containing standard fly media. The fly line containing both Collagen-GAL4 (gift from C. Dearolf) and UAS-GPI::EGFP (gift from S. Eaton) was generated in the lab. Class I and class II myosin RNAi lines were obtained from VDRC and crossed with the generated transgenic GPI::EGFP line. Hemocytes were dissected from 6 third-instar larvae from each cross as previously described(*68*). Briefly, third instar larvae were surface sterilized, and hemolymph was collected by puncturing the integument using dissection forceps into 150 ml of M1G (adjusted to pH=6.9 and additionally supplemented with 1 mg/ml BSA). The medium with cells was resuspended and the cells were allowed to spread for 1-2 hours on a glass bottom dish. Emission anisotropy imaging of the GPI::EGFP was performed on a TIRF microscope and data analyzed as described.

#### siRNA targeting of class I non-muscle myosin

U2OSGG were set to deplete class I myosin proteins. For single isoform knockdown U2OSGG cells were cultured in 6-well plates and transfected using DharmaFect with siRNA-pools against MYO1B (25 nM), MYO1C (25 nM) or MYO1D (25 nM) for 48h to achieve >75% knockdown of the each of the targeted isoforms. To knock down all three isoforms, 25 nM of each of siRNA-pools against MYO1B, MYO1C and MYO1D were co-transfected. Co-transfections were performed thrice over two passages to achieve maximal knockdown of all three isoforms. Transfected cells at each time point were trypsinized and plated for imaging and/or harvested for protein analysis. Data in Supplementary Fig. S2F-J corresponds to anisotropy experiments and immunoblots from cells at the third passage. Non-targeting controls supplied along with the siRNAs were used for every experiment at similar concentrations.

#### Immunoblotting

U2OSGG cells were harvested after each transfection period. Protein lysates were prepared by homogenizing cells in RIPA buffer with protease inhibitors. Protein content was estimated by a BCA estimation kit as per manufacturer’s protocol. 15-20 μg protein was loaded onto 10% Tris-Glycine polyacrylamide gels. Post transfer, PVDF membranes were blocked with Everyblot blocking buffer followed by overnight incubation with primary antibodies, washed and secondary antibody incubation for 1 hour at RT. Membranes were probed with Clarity Max ECL reagent. The expression of class I Myosin proteins was normalized to Tubulin loading controls in the same lanes. Normalization with another housekeeping control, GAPDH, yielded similar expression levels.

#### Bleb cross-linking and analysis

CHO cells (TrVb line, (*69*) stably expressing folate receptor (FR-GPI) or a transmembrane actin binding variant (FRTM-Ez-AFBD*) were transfected with GFP-Myo1c, GFP-Myo1b or LactC2-GFP for 12-15 hours. Blebs were then generated on these cells by incubating with 14 μM Jasplakinolide at 37°C for 30 minutes in M1G. The cells were then incubated with MOV 18(*70*) primary antibody against the folate receptor at 5 μg/mL for 1 hour on ice. This was followed by incubation with 2 μg/mL secondary antibody for 30 min on ice. Samples were imaged immediately after the washing steps taking care to avoid washing away of the weakly cell-attached blebs. Images were obtained on a Zeiss LSM 780 confocal microscope at room temperature using a 63X 1.4NA oil objective by sequential imaging of the cross-linked antibody in far red (647 nm excitation) and the transfected protein in green (488 nm excitation). The blebs were analyzed using a custom MatLab routine that fits the diffraction limited bleb contour, followed by obtaining the average intensity profile along the 5-pixel broadened bleb circumference of both channels. The signal from both channels was cross-correlated, giving a single Pearson’s coefficient for each bleb (Fig 4C).

#### Multi-wavelength emission anisotropy correlation: nanoscale and mesoscale

To correlate emission anisotropy from two distinct membrane components sequential image acquisition was conducted by changing the excitation wavelength and the emission filter. Images were collected as fast as the microscopy system could switch excitation and emission filters ∼1.0 sec.

For the nanoscale analysis, relating anisotropy of one component to the relative local anisotropy of a second membrane component, square ROIs with vertices of 2.1 μm length were selected as indicated above. Only those ROIs where the *I_T_* versus anisotropy was free from noise and trivial-FRET were taking further for analysis (*30*). Next the total intensity of the other channel was determined and used to set a cutoff intensity for the second channel while reporting on the anisotropy of the first channel (see Fig. 4E,F and Supplementary Fig. S3B). Alternatively, it was used to calculate the local intensity ratio, 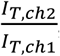, and plotted against the anisotropy calculated from the first channel *r_ch_*_1_ (Fig. 5A).

For the mesoscale image analysis, anisotropy values of both channels were analyzed using a Threshold Overlap Score matrix described previously with minor modifications related to the use of anisotropy values as the scaling factor to select pixels (TOS analysis(*43*)). Briefly, the weighted overlap between pixels of two different channels was calculated depending on the anisotropy threshold for each of the channels. To select comparable thresholds between experiments and cells the distribution of pixel anisotropy values of both channels from each cell was centered around 0 and normalized by its standard deviation following: 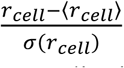, with 〈*r_cell_*〉 being the mean anisotropy and *σ*(*r_cell_*) the standard deviation of the anisotropy distribution from each cell (Fig. 5B and Supplementary Fig. S4A,B). The threshold, *T*, for each channel was next defined as pixels containing a cluster-rich (*T* < 〈*r*〉 − *n* · *σ*(*r*)) or cluster-poor (*T* > 〈*r*〉 + *n* · *σ*(*r*)) organization of the labeled component. The parameter *n* sets the range of values which we defined going from 0 to 3, that is, reducing the selected pixels from the inclusion of all pixels with anisotropy values above or below the average anisotropy to only the pixels that are at least 3 standard deviations away from the mean (see also colored arrows in Supplementary Fig S4B). Each threshold-setting determined the fraction of pixels from the entire cell that were selected for each channel, for channel 1 this would be: 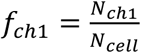, where *N_ch_*_1_ corresponded to the number of channel 1 pixels selected using the threshold and *N_cell_* the total number of pixels associated to the ROI/cell. Overlapping pixels, that are selected for both channels (*N*_ch1=1&2_ see also Fig 5C-E, 6E, F and Supplementary Fig. S4C), combine into a fraction for channel 1 defined as: 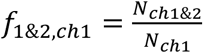.

The ratio with respect to the expected fraction, 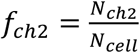 therefore provides a measure for a less than expected (<1) or more than expected (>1) overlap of the thresholded pixels in both channels.

To linearly rescale the average of the two calculated area-overlap ratios, 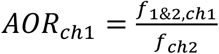, they are corrected for their minimum and maximum values to finally obtain a TOS value that lies between -1 and +1. Here an *AOR* of 1 corresponds to a TOS value of 0. This would mean that in this situation the number of selected pixels from channel 2 overlaps with channel 1 as expected from their area fraction in the cell. For *AOR* < 1 the ratio is rescaled between -1 and 0 through: 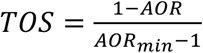. When the number of selected pixels from both channels does not exceed the total number of pixels in the cell, *N_ch_*_1_ + *N_ch_*_2_ < *N_cell_*, none of the pixels have to overlap and this results in a minimum overlap *AOR_min_*_-_ = 0. For *AOR* > 1 the ratio is rescaled between 0 and +1 through: 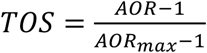. The maximum overlap is dictated by the minimum number of pixels from one of the channels fully overlapping with the other: 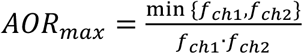. Repeating the above calculations for multiple thresholds in both channels provides a TOS matrix for each cell (Supplementary Fig. S4D). Absence of pixels at a threshold resulted in a TOS value displayed as 0 (Supplementary Fig S4D), but not taken forward into calculating the average TOS matrix values. The image sequence was repeated a second time during each acquisition. This results in a second anisotropy image for each channel that is now shifted in time (about 2 seconds) and functions as a quality control for the acquisition. Assuming minimal reorganization of the myosin activity during this time frame, the TOS matrix between these two images must be the maximally obtainable TOS (Supplementary Fig. S4D,E). Cells with no positive TOS between the anisotropy-low regions (or anisotropy-high regions) of either of the channels were not taken forward for analysis. After this quality control, a single TOS matrix per experiment was generated by averaging the TOS matrices from multiple cells (Supplementary Fig. S4E,F). Day-to-day variability was non-significant for any of the four quadrants of the TOS matrix (Supplementary Fig. S4G).

Simulations of randomly placed pixels chosen from a gaussian distribution of anisotropy values resulted in a TOS matrix that was approximately 0 for all thresholds (Supplementary Fig. S4H). Randomization of the pixels from the experimentally obtained data also resulted in TOS values close to 0, however, the regions with a low number of pixels (the four edges; the most stringent thresholds) contained the largest standard deviation in TOS-values (Supplementary Fig. S4I) and typically had an underestimated TOS-value. Quantification of TOS was therefore obtained from the average values of the bottom left quadrant of the TOS matrix (overlap between low-anisotropy regions) excluding the most stringent four bins (dashed area in the TOS matrix of Supplementary Fig S4E, F).

### Cell spreading experiment and analysis

Fibronectin-coated glass dishes were prepared as mentioned earlier. The cells were de-adhered with TrypLE, pelleted down, re-suspended in incomplete media and left for 30 mins at 37°C for recovery. Next the cells were treated by supplementing the suspensions with the appropriate amount to obtain a final concentration of 0.1% DMSO, 50 µM Blebbistatin or 5 µM PClP and left at 37°C for 60 min. Just before the experiment, the hFN coated dishes containing the appropriate solution were placed in a pre-heated microscopy chamber at 37 °C (OKO labs). Images at 0.1 Hz framerate were recorded using Interference Reflective Microscopy upon adding the pre-treated cells to the dish. Each identified cell in the image-series was analyzed separately using an available MatLab-based software that determined the cell contour at each frame(*71*). The resultant Area-versus-time curves allowed for the calculation of the spreading lagtime, spreading rate and final spread area.

### Interference Reflection Microscopy

Using a 10/90 (reflection/transmission mirror, Thorlabs) unpolarized light filtered at 480/20 (Chroma) from an LED source (pE-400, CoolLED) was directed to the sample using a 60x 1.49NA objective. Reflected light was collected through the same objective and imaged with a sCMOS camera. Through destructive interference of the reflected light, adhering cells were detected as regions of reduced intensity on the camera.

## Acknowledgments

We thank Ram A Vishwakarma (CSIR, N. Delhi) for his support and guidance in chemical synthesis, and Darius Koester (University of Warwick), and all the members of the Mayor laboratory, in particular Chandrima Patra and Sankarshan Talluri for suggestions and feedback on this manuscript. We thank the NCBS Central Imaging and Flow Facility as well as Greeshma Pradeep S for help with optical setups, Pooja Krishna for technical assistance, and computational facilities at NCBS. PS, SS and AB acknowledge doctoral fellowship support from National Centre for Biological Sciences, Tata Institute for fundamental Research (NCBS-TIFR). AB acknowledges the Dedalus community (Google group) and free access to Dedalus provided by the developers.

## Author contributions

Conceptualization: PS, TSvZ, SM. Methodology: PS, TSvZ, SS, SJ, AB, MK, PPS, MR, SM. Investigation: PS, TSvZ, SS, SJ, AB, BM, MK. Visualization: PS, TSvZ. Funding acquisition: SM. Project administration: PPS, MR, SM. Supervision: TSvZ, MR, SM. Writing – original draft: PS, TSvZ, MR, SM. Writing – review & editing: PS, TvZ, SJ, AB, BM, MK, PPS, MR, SM

## Data and materials availability

All data are included in the figures in main text or the supplementary materials.

## Supplementary Materials

Supplementary Text Supplementary Figs. S1 to S6 Supplementary Tables S1 to S5 References

