## Supplementary Material for "Diverse ancestral myosin motors generate and segregate distinct types of nanocluster-rich domains at the plasma membrane"

(next symbols should be: #, **, ††, ‡‡, etc.)

**The Supplementary Materials PDF file includes:**

Supplementary Text
Figs S1 to S6
Tables S1 to S5
References (72-80)

**Supplementary Text**

Active Flory-Huggins theory for a three-component membrane coupled to the actomyosin cortex:

In previous studies(*26*, *72*), we developed an Active Flory-Huggins (AFH) theory for a multicomponent membrane interacting with the actomyosin cortex comprising a single myosin species. Here we extend our framework to include two myosin species, namely the class I and class II myosin, with differential turnover rates and contractility.

Consider a planar membrane, composed of a ternary mixture of saturated lipids (*lo* component, that includes GPI-APs), unsaturated lipids (*ld* component) and transmembrane proteins (TM-EzAFBD), with area fraction $\phi_1$, $\phi_2$ and $\psi$ respectively, such that $\phi_1 + \phi_2 + \psi = 1$. At temperature above $T_c$, the membrane is in a homogeneous, mixed phase. At temperatures below $T_c$, the mixture undergoes an *lo-ld* equilibrium phase separation, with the saturated lipids predominantly in the liquid-ordered (*lo*) phase and the unsaturated lipids in the liquid-disordered (*ld*) phase.

On coupling to the actomyosin cortex, the *lo* component experiences a force from active stresses generated through class I non-muscle myosin and the direct actin binder TM-EzAFBD experiences a force from the active stresses generated through class II non-muscle myosin. The dynamics of the area fractions of the components follow from local force balance and mass conservation (*26*, *72–74*),

$$\frac{\partial \phi_1}{\partial t} = \nabla \cdot [-L\left(\nabla \mu_\psi - K_{II}\,\nabla \cdot \sigma_{II}\right) + M_1(\nabla \mu_1 - K_I\,\nabla \cdot \sigma_I)]$$

$$\frac{\partial \psi}{\partial t} = \nabla \cdot [-L\,(\nabla \mu_1 - K_I\,\nabla \cdot \sigma_I) + M_\psi\left(\nabla \mu_\psi - K_{II}\,\nabla \cdot \sigma_{II}\right)] \qquad (1)$$

where the exchange mobilities(*73*, *74*) are given by $L \propto \phi_1\,\psi$, $M_1 \propto \phi_1(1-\phi_1)$, and $M_\psi \propto \psi(1-\psi)$, and the currents in the right-hand side of the above equations have contributions from the equilibrium exchange chemical potentials $\mu_1$ and $\mu_\psi$, and the nonequilibrium active contractile stresses $\sigma_I$ and $\sigma_{II}$, arising from the action of class I and class II myosin, respectively. The exchange chemical potential is obtained from the Flory-Huggins free energy $F$,(*72*) specifically,

$$\mu_1 = \chi_{12}(\phi_2 - \phi_1) + \left(\chi_{1\psi} - \chi_{2\psi}\right)\psi + \ln(\phi_1/\phi_2) - \kappa\nabla^2(\phi_1 - \phi_2)$$

$$\mu_\psi = \chi_{2\psi}(\phi_2 - \psi) + \left(\chi_{1\psi} - \chi_{12}\right)\phi_1 + \ln(\psi/\phi_2) - \kappa\nabla^2(\psi - \phi_2) \qquad (2)$$

where $\chi_{12}$, $\chi_{1\psi}$, $\chi_{2\psi}$ are the Flory-Huggins interaction parameters between the membrane components controlling affinity of *lo* with *ld*, *lo* with TM-EzAFBD, and *ld* with TM-EzAFBD, respectively.

The parameters $K_I$ and $K_{II}$ characterise the coupling between class I myosin with *lo* component and class II myosin with TM-EzAFBD, respectively. The active cortical stresses $\sigma_{I,II}$ in equation (1) are evaluated from the active hydrodynamic equations describing the actomyosin cortex (*72*) in terms of the local concentration of actomyosin (equations for mass balance) and the cortical velocities of bound class I and class II myosins (equations for force balance). The form of the active stresses, exerted by bound class I and class II myosins (Supplementary Fig S5A, D) is given by:

$$\sigma_I = \left(\frac{\zeta_I\, c_I}{1 + c_I} - Q_I c_I c_{II}\right) I$$

$$\sigma_{II} = \left(\frac{\zeta_{II}\, c_{II}}{1 + c_{II}} - Q_{II} c_I c_{II}\right) I$$

(3)

where in the absence of any long range orientational order, the active stresses are isotropic (thus in the above equation, $I$ is the identity matrix). The first term shows the usual saturating form with $\zeta_I, \zeta_{II} > 0$ for contractile stresses (*75*). The second term represents a potential inhibition of the local activity of bound myosin motor I due to the presence of myosin motor II (and *vice versa*). While in general $Q_I$, $Q_{II}$ can have different values, we have, for convenience, taken them to be the same and equal to $Q$.

The mass balance equations express the balance between advection and turnover of the bound myosin species. The turnover of myosin has contributions from unbinding and binding. Here, we take the unbinding rate to be constant and choose the following simple form for the binding rates,

$$k_{bI}(\phi_1, \psi, c_I, c_{II}) = A_I \phi_1 - B_I c_{II}$$

$$k_{bII}(\phi_1, \psi, c_I, c_{II}) = A_{II} \psi - B_{II} c_I \qquad (4)$$

which includes specific binding $A_I, A_{II}$ to the membrane components, *lo* and TM-EzAFBD, respectively, thereby generating positive feedback, and competitive binding $B_I, B_{II}$ between the myosin species.

This completes our description of the active Flory-Huggins (AFH) theory, for the three-component membrane coupled to an active cortex comprising two myosin species. The explicit form of the active hydrodynamic equations of the actomyosin cortex are presented in (*72*). Using this, we study the steady state configurations of the membrane composition at a fixed temperature below and above $T_c$ starting from mixed initial condition. As seen in (*26*, *72*) even at temperatures above $T_c$, we observe mesoscale segregation of the *lo* and *ld* components, arising from active stresses and lateral interactions between the lipid species. Even though TM-EzAFBD has no affinity to either *lo* or *ld* components, it segregates from the *lo* components (Fig S5G and Supplementary Fig S5B,C) due to differential active stresses generated by myosin I and myosin II. The extent of overlap between the *lo* and TM-EzAFBD domains depends on the Flory interaction parameters $\chi_{1\psi}$, $\chi_{2\psi}$ (Fig 5G,H and Supplementary Fig S5E,F) and the actomyosin parameters $B_I, B_{II}$, and $Q$ (Fig 5H). As follows clearly from our discussion here and in the main text, the effect of changing these parameters on the segregation of the *lo* component and TM-EzAFBD domains is displayed in Supplementary Table 5 below. A striking observation is that the domains of *lo* and TM-EzAFBD exhibit strong density inhomogeneities which we refer to as *granularity* (see main text).

Supplementary Figure 1 Photobleaching

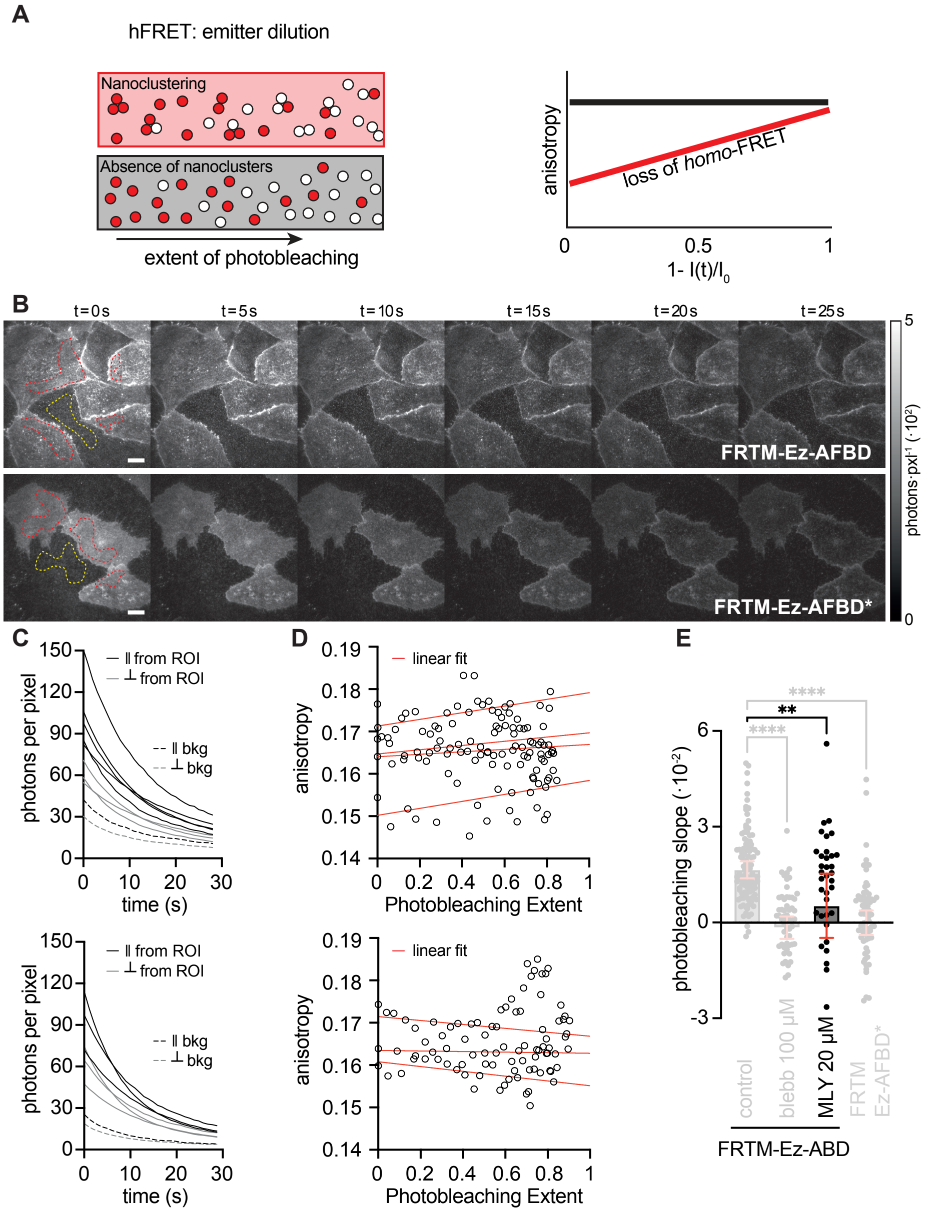


**Supplementary Figure S1. Photo-dilution experiments on FR-labeled cells and workflow to extract the anisotropy-photodilution slope. A)** Cartoons (left) provide a pictorial representation of molecules (red circles) organized as monomers with nanoclusters (pink background) or monomers only (grey background) during photobleaching, which results in the dilution of fluorophores. Representative plot (right) of the change in fluorescence emission anisotropy of the monomers with nanoclusters (red) and monomer only (black) panels following photobleaching. At the start of the photobleaching process, the collection of monomers and nanoclusters (pink panel) have a lower collective anisotropy due to homo-FRET between fluorophores in nanoclusters compared to those that are present only as monomers (grey panel). Following photobleaching the progressive loss of fluorophores (indicated as open circles) results in only monomers remaining in both cases, and the emission anisotropy values equalize. The slopes of these plots provide a method to compare the extent of nanoclustering. **B)** Montages of total fluorescence intensity images (calculated through ⊥ and ∥ channels) from PLB-labelled FRTM-Ez-AFBD and FRTM-Ez-AFBD* show the loss of fluorescence as a function of progressive photobleaching. Regions of interest from cells are indicated by a red-dashed lines and the background region is selected with the yellow-dashed line. **C)** The traces from PLB-labelled FRTM-Ez-AFBD (top) and FRTM-Ez-AFBD* (bottom) cells capture emission intensity decays of the orthogonal polarisations from specific regions of interest indicated in panel B. **D)** Plots depict the change in emission anisotropy (y-axis) as a function of the extent of photobleaching (x-axis). Each region of interest is plotted as a scatter plot and fitted to linear equations (red lines; $y = mx + c$). **E)** Bar graph shows the mean (±s.d.) of the slopes extracted from the linear fits of the data derived from the FRTM-Ez-AFBD construct treated with MLY (38 cells, N=2). The grey bars depict data already shown in Fig. 2C to allow direct comparison. Scale bars, 10 μm. ****, and ** correspond to $p$-values of $<10^{-4}$, and $<10^{-2}$, respectively.

Supplementary Figure 2 Myosin 1 siRNA

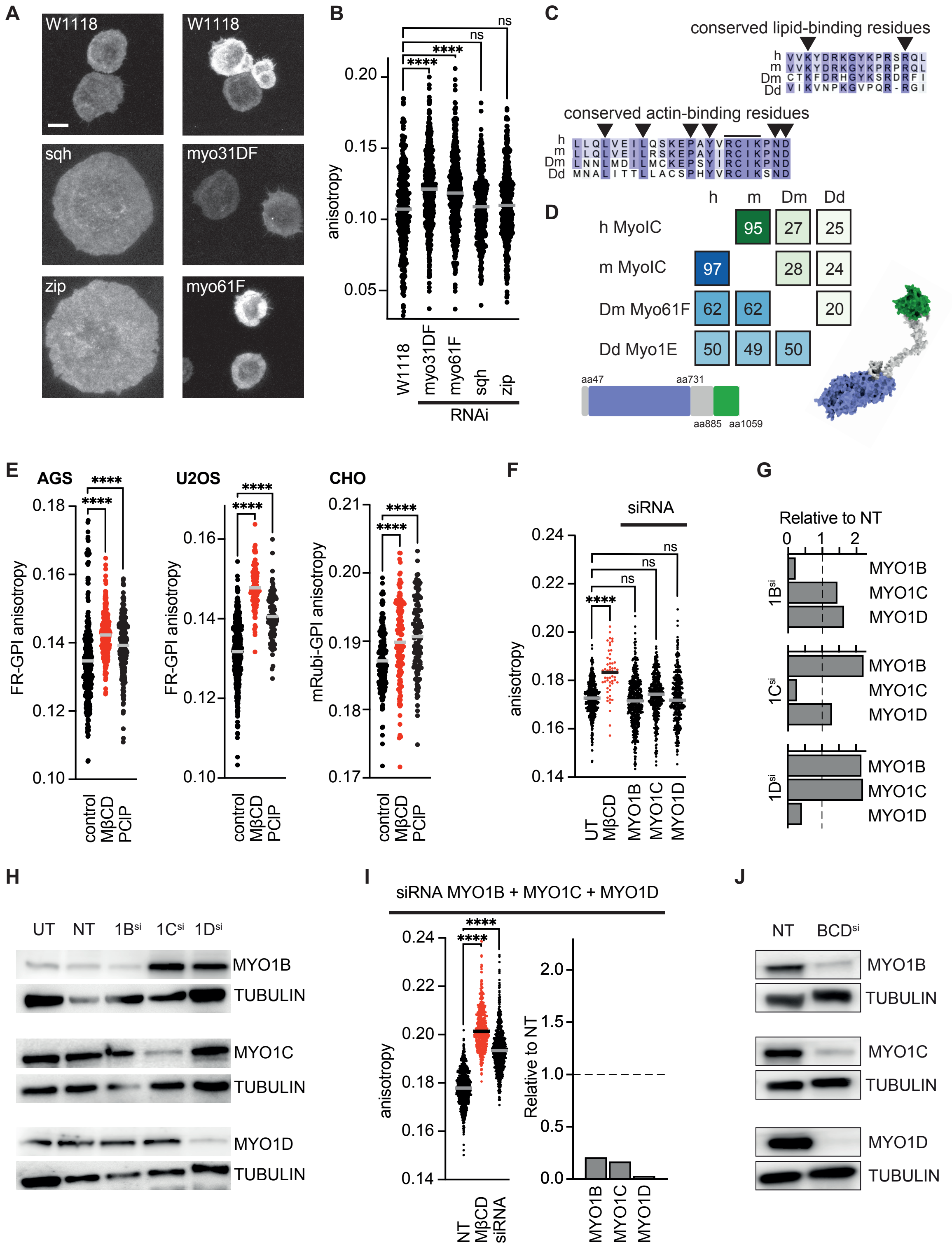

**Supplementary Figure S2. Myosin class I siRNA knock down. A)** Montages of total fluorescence intensity images (calculated through $\boldsymbol{I}_{\parallel}$ and $\boldsymbol{I}_{\perp}$ channels) of hemocytes derived from a *D. melanogaster* GPI::GFP transgenic crossing with a transgenic containing the indicated UAS-dsRNAi. **B)** Graphs showing anisotropy values from the corresponding GFP-GPI expressing hemocytes expressing the indicated dsRNAi. The data set was obtained from 55 cells (340 ROIs), 83 cells (477 ROIs), 77 cells (467 ROIs), 33 cells (348 ROIs), and 34 cells (452 ROIs) for the cross with W1118, myo31DF, myo61F, sqh, and zip, respectively. **C,D)** Conserved residues of the lipid binding (green shades) and actin binding domains (blue shades) of Myosin1c homologs across, mammals (human, mouse, fruitflies (Drosophila) and protists (Dictyostelium) (i) and percentage sequence homology between Myo 1c proteins across these species (j). **E)** Graphs showing anisotropy values from cells treated with the vehicle (DMSO), MβCD (10 mM), or myosin 1 inhibitor (PClP at 5 μM). The data is collected from FR-GPI, labeled with PLB, and expressed in AGS cells (left graph: 21 cells {307 ROIs}, 28 cells {333 ROIs}, and 23 cells {280 ROIs} for control, MCD and PClP, respectively), FR-GPI expressing U2OS cells (middle graph: 23 cells {414 ROIs}, 9 cells {116 ROIs}, and 26 cells {91 ROIs} for control, MCD and PClP, respectively) and mRubi-GPI expressing CHO cells (right graph: 17 cells {177 ROIs}, 16 cells {143 ROIs}, and 20 cells {129 ROIs} for control, MCD and PClP, respectively). **F)** Anisotropy distributions of GFP-GPI in U2OS cells depleted of Myosin1 isoforms. Graph shows anisotropy values of cells that were treated with siRNA against Myo1B, Myo1C, or Myo1D. Anisotropy values of the siRNA treatments and cells treated with 10 mM MβCD to abrogate GFP-GPI clustering were compared to the untreated control (UT). The anisotropy data was compiled from 47 cells (468 ROIs), 27 cells (56 ROIs), 44 cells (484 ROIs), 29 cells (383 ROIs), and 29 cells (351 ROIs) for the untreated cells, MβCD, MYO1B siRNA, MYO1C siRNA, and MYO1D siRNA, respectively. **G)** Bar graph shows levels of MYO1B, C&D protein expression in U2OS cells that were treated with siRNAi against Myo1B, Myo1C, or Myo1D. Protein levels for Myo1B, Myo1C and Myo1D were normalized to the non-target control. **H)** Representative western blot showing the reduction of the respective myosin protein bands with the loading controls for the untreated cells, the non-target siRNA and the three individual myosin 1 siRNA. **I)** Graph shows anisotropy values of cells that were treated with a combined cocktail of siRNA against Myo1B/Myo1C/Myo1D compared to the non-target control and the 10 mM MβCD. The data was compiled from 40 cells (666 ROIs), 29 cells (456 ROIs), and 64 cells (711 ROIs) for the non-target siRNA, MβCD, and triple siRNA knock downs, respectively. Protein levels for Myo1B, Myo1C and Myo1D were also compared to the non-target control. Note that only when a triple siRNA is used, there is a significant increase in emission anisotropy comparable to that seen with the loss of clustering due to MβCD-treatment. **J)** Representative western blot showing the reduction of the respective myosin protein bands with the indicated loading controls. Scale bar, 10 μm. ****, and ns correspond to *p*-values of $<10^{-4}$, and >0.05, respectively.

Supplementary Figure 3: mCherry-LactC2-EzAFBD expression in cells

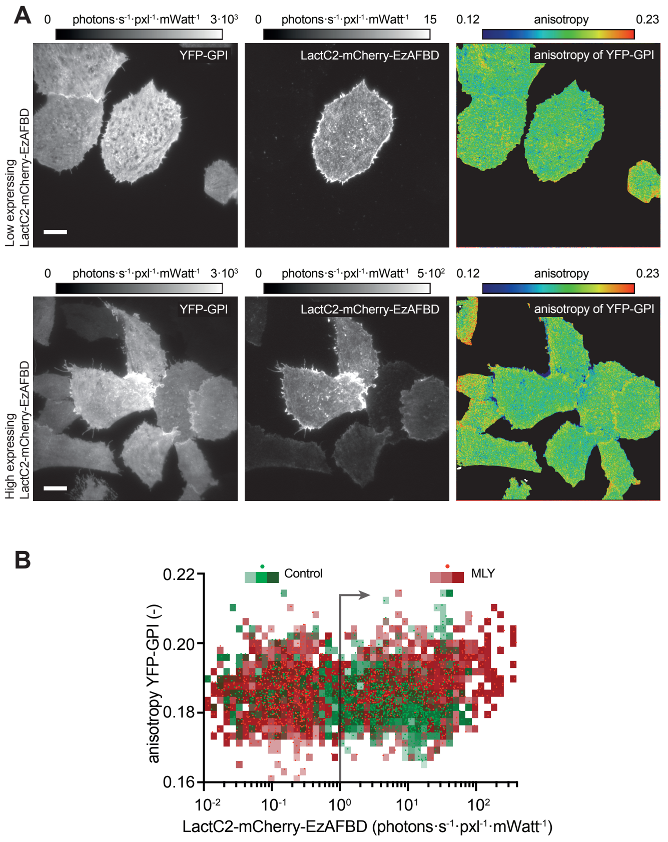


**Supplementary Figure S3. Co-expression of a synthetic PS-actin linker protein and GPI-AP. A**) Intensity and anisotropy (right panel) images of cells co-expressing YFP-GPI (left panels) and different levels of the PS-actin linker, mCherry-LactC2-EzAFBD (center panel). Intensity images display the total photon counts from each fluorophore. **B**) Scatter plot shows anisotropy of YFP-GPI as a function of the (local) expression level of the mCherry-LactC2-EzAFBD chimeric protein. In green are the ROIs from control cells and in red the ROIs from cells pre-treated with the MLY cocktail. The line and arrow indicate the threshold intensity levels applied to distinguish cells that express the linker and cells that do not. Each data point represents an ROI measurement, and the squares are binned values of expression level and anisotropy, false colored for the control (green) and MLY-treated (red). **C**) Intensity (left and middle) and anisotropy (right) images of YFP-GPI cells, co-expressing mCherry-LacC2-EzAFBD (middle) exhibiting sensitivity to the MLY cocktail. Scale bars for A and C, 10 µm.

Supplementary Figure 4 Threshold Overlap Score analysis

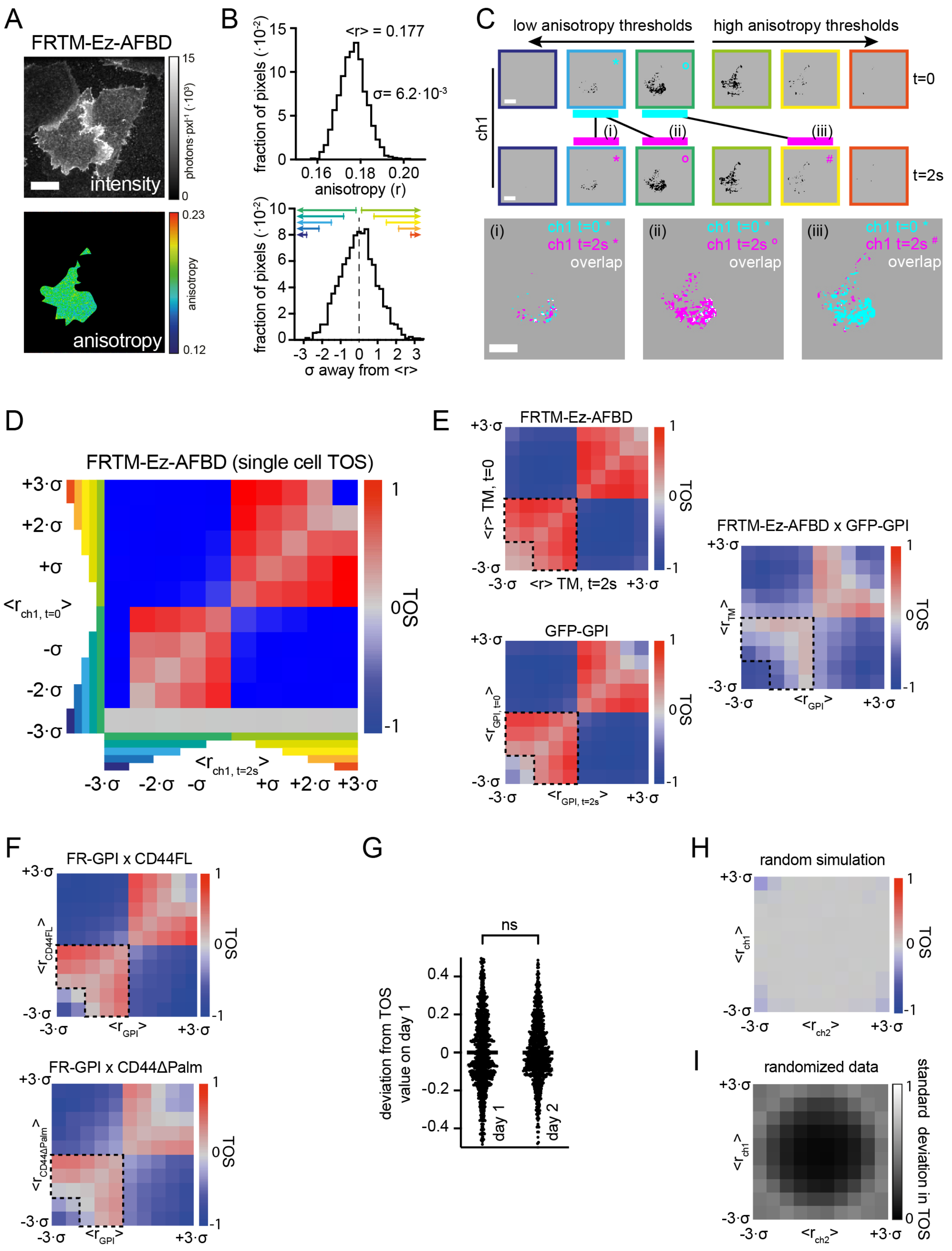

**Supplementary Figure S4. Threshold Overlap Score (TOS) analysis. A**) Total intensity image (top panel) and anisotropy image (bottom panel). The total intensity image shows the entire field of view whereas the anisotropy image displays the selection of the cell of interest that is taken forward for analysis. **B**) The distribution of anisotropy values from the selected cell from a (top panel). The distribution is shifted with respect to the mean anisotropy value and normalized to the standard deviation of the original distribution (bottom panel). **C**) Example thresholded images showing selected pixels (black on a gray background) from the same channel at the two different timepoints (top and bottom row). The designated threshold values from low to high anisotropy regions (left to right: colored frames are correlated to colored arrows in B) result in different sets of pixels being selected. Examples of overlap between designated thresholds for the two timepoints of FRTM-Ez-AFBD in the same cell are displayed in panels i-iii. **D**) TOS-matrix of the cell selected in a resulting from the calculation of the TOS-values for all the threshold combinations. **E**) Averaged TOS matrix from multiple cells (58 cells, N=2) from temporal FRTM-Ez-AFBD correlation, temporal GPI-AP correlations and the FRTM-Ez-AFBD with GPI-AP cross-correlation. **F**) Average TOS matrix for FR-GPI with CD44FL-GFP (52 cells, N=2) and FR-GPI with CD44ΔPalm-GFP (48 cells, N=2). The area enclosed by the dashed lines in e and f was used to calculate the mean TOS-values per cell reported in Fig 5**E** and **L**. **G**) Day to day variability of mean TOS-values normalized to day 1 (day 1: 1020 values from 255 cells, and day 2: 876 values from 219 cells) from any of the four quadrants in a TOS matrix. **H**) TOS-matrix from simulations of random spatially distributed anisotropy values. **I**) Matrix of the standard deviation in TOS-values derived from a randomized distribution of the data used for Fig. 5E,L and Supplementary Fig 5E,F. Scale bars, 10 μm for A and C. ns correspond to *p*-values of >0.05.

Supplementary Figure 5 Numerical Simulations Active Flory-Huggins Theory

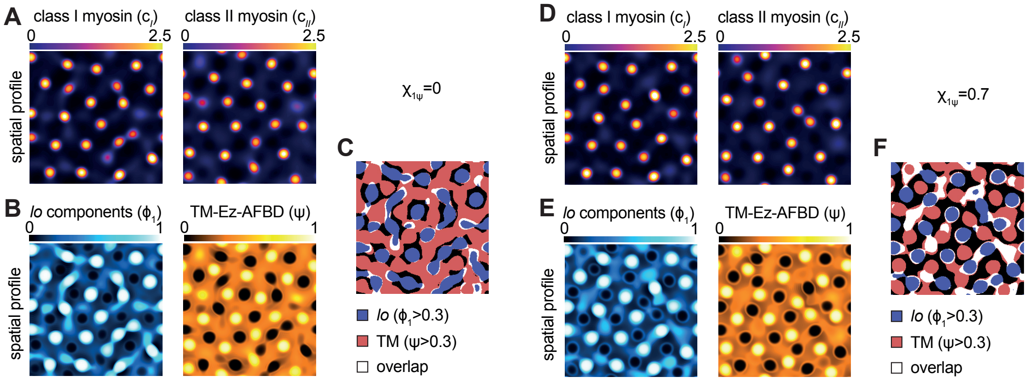


**Supplementary Figure S5. Numerical simulations for active Flory-Huggin theory. A and D)** Concentrations of the active myosin species: class I myosin ($c_1$; left) and class II myosin ($c_2$;

right). **B and E)** Area fractions of the *lo* phase ($\Phi_1$; left) and the TM-Ez-AFBD ($\Psi$; right) due to their interaction with the active stresses generated by the myosin distribution displayed in a and d. **C and F)** Mesoscale domain overlap, where domains are identified based on their area fractions >0.3. Simulation corresponding to a-c used the interaction parameter $\chi_{1\psi} = 0$ and the simulations corresponding to d-f used $\chi_{1\psi} = 0.7$.

Supplementary Figure 6: Cell spreading

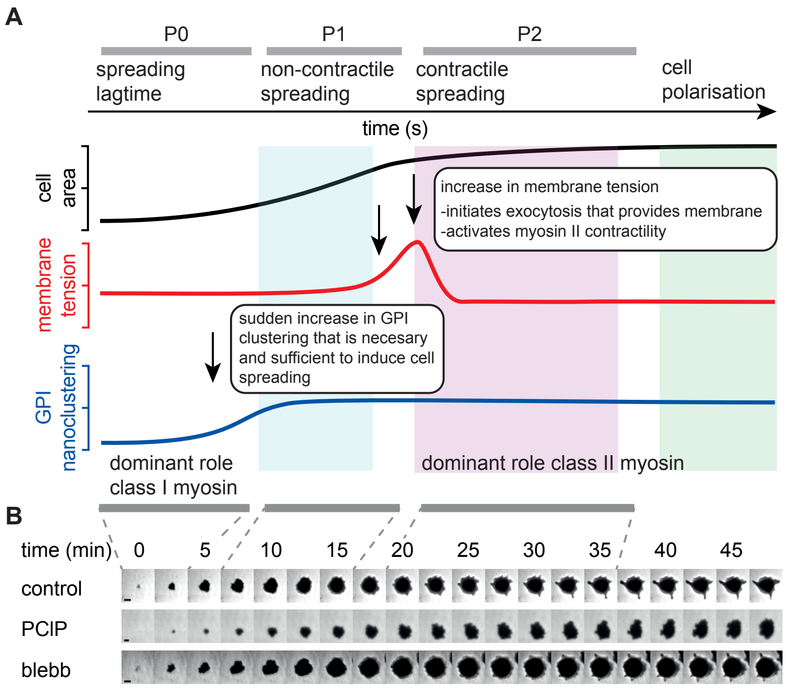


**Supplementary Figure S6. Functional consequences of myosin class independence on membrane patterning on cell spreading. A)** Schematic showing the different phases of cell spreading, their relation to GPI-AP nanoclustering and membrane tension changes following published studies and the periods of expected contributions from class I and class II myosin motor activity(*49–51*). **B)** Image montages (top) and temporal evolution of spread area (bottom) of cells treated with vehicle (DMSO, 1 hour), PCIP (5 µM, 1 hour) or Blebbistatin (blebb 50 µM, 1 hour) and adhering to fibronectin-coated surfaces imaged using Interference Reflection Microscopy. **C-E)** Graphs showing the time-dependent cell spread area per cell (C), time lag between cell attachment and cell spreading (D) and the extent of cell area increase (E; mean ± s.d) during spreading of the same cells as in C where the maximum rate of area increase was shifted to t=0. These data have been obtained from 11, 14 and 19 cells for DMSO, blebb and PClP, respectively, N=3. Scale bars, 10 µm. **, and ns correspond to *p*-values of $<10^{-2}$, and >0.05, respectively.

### Supplementary Table 1. List of constructs

| Construct name | Description | Vector / primer / Restriction site / Resistance | Source |
|---|---|---|---|
| CD44-GFP | Mouse CD44 standard isoform tagged to GFP | pEGFPN1; Kan, BglII, EcoRI sites | (*76*) |
| CD44ΔPalm-GFP | Mouse CD44(C290A,C299A) defective in palmitoylation | pEGFPN1; Kan, BglII, EcoRI sites | (*28*) |
| EGFP-GPI(M3GG) | EGFP tagged to the GPI signal of the folate receptor | pJB20; Amp | (*77*) |
| FR-GPI | Human folate receptor | pMEP4; Amp | (*34*) |
| YFP-GPI | YFP tagged to GPI anchor | pEGFP-N1(GFP replaced by YFP); Kan | (*31*) |
| mRuby GPI | EGFP replaced by mRuby in M3GG | pJB20; Amp | (*51*) |
| mCherry-LactC2-EzAFBBD | mCherry-LactC2 (PS binding domain) fused with actin binding domain of Ezrin | pcDNA3.1, Amp, cloned using Gibson assembly | In this study |
| FRTM-Ez-AFBD | Folate Receptor tagged to TM-domain of IgG and AFBD of Ezrin | pcDNA3.1; Amp; HindIII and NotI | (*14*) |
| FRTM-Ez-AFBD* | Single point mutation in AFBD domain of the FRTM-Ez-AFBD proteins to reduce actin binding | pcDNA3.1; Amp; HindIII and NotI | (*14*) |
| GFP-Myo1c | Wildtype Myosin1c isoform (mouse) | pEGFP-N1, Kan | (*78*) |
| GFP-Myo1b | Wildtype Myosin1b isoform (mouse) | Kan | (*79*) |
| mCherry-Myo2a | Wild type non-muscle Myosin2a tagged to mCherry | Kan | Gift from Mike Sheetz |

### Supplementary Table 2. List of Cell Lines

| Cell lines | Description | Source |
|---|---|---|
| GG8 | CHO-K1 cells (TRVb-1), devoid of transferrin receptor (TfR) and stably transfected with human TfR and EGFP-GPI | (*69*) |
| FR-GPI | CHO-K1 cells (TRVb-1), devoid of transferrin receptor (TfR) and stably transfected with human TfR and FR-GPI | (*31*) |
| MYG1; m-YFP-GPI expressing CHO cells | CHO-K1 cells (TRVb-1), devoid of transferrin receptor (TfR) and stably transfected with human TfR and YFP-GPI | (*51*) |
| CHO-FRTM-Ez-AFBD | CHO-K1 cells (TRVb-1), devoid of transferrin receptor (TfR) and stably transfected with human TfR and EGFP-GPI | (*14*) |
| CHO-FRTM-Ez-AFBD* | CHO-K1 cells (TRVb-1), devoid of transferrin receptor (TfR) and stably transfected with human TfR and EGFP-GPI | (*14*) |
| FR-AGS | Human AGS cells stably transfected with FR-GPI | (*14*) |
| U2OS-GG | YFP tagged to GPI anchor | (*51*) |

### Supplementary Table 3. List of fly-lines

| Name | Description | Source |
|---|---|---|
| w[1118] | control | Bloomington |
| UAS GPI::GFP | Gift from Susan Eaton | (*80*) |
| Gal4 collagen | Gift from C Dearolf | (*80*) |
| w[1118];P{GD1695}v7916 | sqh (CG3595) dsRNAi under UAS promotor | VRDC |
| w[1118];P{GD1566}v7819 | zip (CG15792) dsRNAi under UAS promotor | VRDC |
| P{KK102456}VIE-260B | myo31DF (CG7438) dsRNAi under UAS promotor | VRDC |
| P{KK101033}VIE-260B | myo61F (CG9155) dsRNAi under UAS promotor | VRDC |

### Supplementary Table 4. List of Reagents

| Name | Source |
|---|---|
| Fetal Bovine Serum (FBS) | Gibco, 16000044 |
| Ham's F12 media | HiMedia AT144-5L |
| penicillin-streptomycin-glutamine cocktail | Sigma, G1146 |
| TrypL Express | Thermo Fisher Scientific 12-604-039 |
| Fugene6 | Promega, E2692 |
| McCoy's 5A | HiMedia AT057A |
| Human Fibronectin | Merck Millipore |
| Helmanex III | Hellma Analytics |
| PLB | Synthesized by PP Singh CSIR-IIIM, Jammu, India |
| DharmaFECT | T-2001-02 |
| ON-TARGETplus Non-targeting Pool | D-001810-10-05 (Dharmacon) |
| MYO1B SMART siRNA pool | L-023110-01-0005 (Dharmacon) |
| MYO1C SMART siRNA pool | L-015121-00-0005 (Dharmacon) |
| MYO1D SMART siRNA pool | L-023316-01-0005 (Dharmacon) |
| MemBrite Fix 405/430 | Biotium, 30092-T |
| Phalloidin Abberior Star 635 | Sigma, 30972 |
| Myosin 1B Antibody | Novus Biologicals, NBP1-87739 |
| Myosin 1C Antibody | Novus Biologicals, NBP1-87745 |
| Myosin 1D (E6TT6Q) antibody | Cell Signaling Technology, 90307s |
| Anti-Beta Tubulin antibody (clone, AA2) | Sigma-Aldrich, T8328 |
| Pierce microBCA protein assay kit and reagents | Thermo Scientific, 23235 |
| cOmplete™, EDTA-free Protease Inhibitor Cocktail | Millipore Sigma, 11873580001 |
| Ripa Buffer (10X) | Cell Signaling Technology, 9806S |
| Immobilon-P PVDF Membrane | Merck, IPVH00010 |
| EveryBlot Blocking Buffer | Bio-Rad, 12010020 |
| Clarity Max™ Western ECL Substrate | Bio-Rad, 1705062 |
| MβCD | Sigma Aldrich, C4555 |
| Jasplakinolide | Invitrogen, J7473 |
| Blebbistatin | Sigma Aldrich, B0560 |
| Y-27632 | Sigma Aldrich, Y0503 |
| ML7 | Sigma Aldrich, I2764 |
| Pentachloropseudilin (PClP) | Synthesized in the lab of PP Singh and Ram V. CSIR-IIIM, Jammu, India (Ref 35) |

### Supplementary Table 5: Qualitative effect of simulation parameters

| Changes in AFH parameters | Physical interpretation of parameters | Effect on segregation of *lo* component and TM-EzAFBD |
|---|---|---|
| Increase in $B_{I,II}$ | Competitive binding between class I and class II myosin | Decreases domain overlap |
| Increase in $Q$ | Competitive inhibition of active stresses | Decreases domain overlap |
| Increase in $\chi_{2\psi}$ | Affinity between *ld* component and TM-EzAFBD | Increases domain overlap |
| Increase in $\chi_{1\psi}$ | Affinity between *lo* component and TM-EzAFBD | Promotes segregation and decreases domain overlap |
| Increase in $K_I,\ K_{II}$ | Characterizes coupling between class I myosin with *lo* component and class II myosin with TM-EzAFBD | Promotes nano- and mesoscale segregation of *lo* components and TM-EzAFBD |
